# Synthetic Biomolecular Condensates as Tunable Microtubule Assembly Hubs

**DOI:** 10.64898/2026.05.01.722010

**Authors:** Sukanya Srinivasan, Anurag Singh, Davit A. Potoyan, Priya R. Banerjee

## Abstract

Phase separation of proteins and nucleic acids (NAs) into nano-to-microscale condensates can regulate biochemical processes, including assembly and organization of cytoskeletal networks such as actin and microtubules. This study examines the functional role of condensate material properties in microtubule assembly. Learning from the sequence grammar of naturally occurring intrinsically disordered regions in microtubule-associated proteins, two-component peptide-NA condensates with programmable material properties were designed. These synthetic condensates catalyze tubulin polymerization into microtubule filaments with tunable outcomes. Tubulin preferentially partitions to the condensate interface and nucleates microtubule assembly. Enhanced tubulin self-assembly produces long filaments that exhibit branching and bundling. Using a minimal stochastic chemo-mechanical model, we show that sequence-encoded condensate viscoelasticity is a tunable element that controls filament morphologies and identifies interfacial rheology as the key regulator of filament growth. Fluorescence recovery after photobleaching experiments support this model, revealing a direct correlation between interfacial tubulin mobility and condensate-directed microtubule assembly. Distinct regimes emerge due to competition between bulk adsorption and lateral diffusion of tubulin at the condensate interface, which determines whether filament tips grow or stall. Since dynamic microtubule assembly and restructuring are essential for various cellular functions, this work highlights a critical role of condensate interfacial rheology in cytoskeletal organization.

## Introduction

Biomolecular condensates are formed via multivalent interactions among proteins, nucleic acids (NAs), and other macromolecules, and can play crucial roles in organizing subcellular processes across diverse biological systems^1, 2^. Many condensate-forming proteins contain intrinsically disordered regions (IDRs) whose amino acid composition and patterning encode conformational heterogeneity ^3^. The “molecular grammar”, defined by the composition and patterning of amino acids in their linear sequence, dictates the strength and valence of intra- and intermolecular interactions, thereby governing partner binding, ensemble dimension^4–6^, condensate formation, and the stability, composition, and material properties of the resulting materials^7–13^.

Condensates can create distinct microenvironments that differ markedly from the surrounding dilute phase, exhibiting localized pH gradients^14–16^, ion partitioning^17–19^, and unique viscoelastic material properties^7, 20–22^ that can collectively influence macromolecular diffusion and reaction dynamics^17, 23, 24^. Additionally, condensate size and morphology can modulate functional output by regulating interfacial properties, molecular partitioning, and internal crowding^25–27^. Together, these features offer tunability in condensate structure and function through multiple, degrees of freedom, opening the door to systematic exploration in synthetic biology applications, especially in the context of protocell research^28^.

Synthetic condensates provide a powerful platform to test these principles and to determine condensate properties that are crucial for their function as crucibles for biochemical reactions. By enriching enzymes and their substrates, condensates can, in principle, enhance biochemical reactions^29, 30^. However, condensates can also dampen reaction rates due to altered molecular conformations, binding affinities, and slow diffusion dynamics within viscoelastic networks^31–33^. Indeed, both acceleration and dampening of ribozyme catalysis have been observed in engineered condensates^17, 34, 35^. Synthetic condensates, therefore, offer a unique opportunity to dissect how condensate-specific physical and chemical grammar regulates biochemical reactions in a context-specific manner.

Protein condensates have recently emerged as regulators of cytoskeletal dynamics, serving as reaction hubs that promote nucleation and growth of cytoskeletal networks that are essential for mitosis, intracellular transport, and cell polarity^36^. Condensates formed by several actin-interacting proteins, as well as those containing components of the T cell receptor phosphorylation cascade, can assemble and bundle actin filaments^37–41^. Similarly, the bacterial tubulin homolog, FtsZ, forms condensates with its interaction partners, from which FtsZ filaments polymerize upon GTP addition^42, 43^. Notably, synthetic condensates have been shown to scaffold actin^44^ and FtsZ, leading to enhanced filament assembly, lateral filament association (bundling), and condensate deformation^45^. In C. elegans, centrosomes provide a compelling *in vivo* case for condensate-mediated microtubule (MT) nucleation and growth^46^. Meanwhile, *in vitro* studies have demonstrated that condensation of the spindle matrix protein ^47^ and various microtubule-associated proteins (MAPs)^48^ can locally enrich tubulin, promote MT nucleation, and spatially coordinate branching MT assembly^49^. Similarly, Tau, a MAP, forms condensates that locally nucleate MT bundles by enriching tubulin. These seminal studies collectively underscore the importance of the condensate microenvironment in this process^50–53^.

Inspired by the lysine-rich MT-binding IDR of Tau and the ability of cationic polypeptides to engage in electrostatically driven coacervation with anionic NAs, we report a bottom-up approach to create synthetic peptide-NA condensates that can function as tunable chemo-mechanical hubs for MT self-assembly. Although studies in C. elegans revealed that regulated changes in centrosome material properties are important for cytoskeletal organization^36, 54, 55^, the precise role of condensate material properties (viscoelasticity, client recruitment, and their mobility) in controlling MT dynamics has yet to be systematically evaluated. By manipulating the chemical and physical properties of peptide and NA components, we can generate programmable condensates with defined viscoelasticity, internal composition, and interfacial properties^21, 22, 56, 57^. These features provide a tunable platform to investigate how mesoscale material properties modulate tubulin partitioning, MT assembly, and other biochemical reactions.

Using time-lapse confocal fluorescence imaging and computational modeling, here we demonstrate that synthetic peptide-NA condensates can selectively partition tubulin and catalyze MT nucleation and growth at the condensate interface. By modulating condensate viscoelasticity and interface properties through peptide sequence design, NA length, and mixture composition, we show that condensate material properties exert differential effects on MT nucleation, dynamics, bundling, and spatial organization. Finally, by developing a minimal, coarse-grained stochastic kinetic model, we provide mechanistic insights into the condensate viscoelasticity-directed regulation of MT assembly dynamics.

## Results

### Synthetic peptide-NA condensates catalyze MT assembly

The sticker-and-spacer model provides a framework for understanding multivalent interactions that drive biomolecular condensation in disordered polypeptides with low complexity sequences^9^. Based on the previous studies on RNA-binding peptides, arginine (R) and tyrosine (Y) residues in IDRs can act as “stickers” that enable inter-chain attractive interactions, while “spacer” residues such as glycine (G) and proline (P) regulate interactions mediated by these stickers as well as the chain solvation properties^21^. To identify unique sequence features that may encode phase separation and/or tubulin and MT binding among MAP IDRs, we performed a bioinformatics analysis of IDRs from >1000 human MAPs by calculating an enrichment score for the fraction of each residue relative to the human proteome. Interestingly, this analysis revealed a compositional bias characterized by enrichment of proline (P), serine (S), and lysine (K), and depletion of hydrophobic and aromatic amino acids (**Figure 1A**). Recently, IDR grammars inferred using NARDINI+ (GIN) highlighted the enrichment of K-blocks in nucleolar proteins involved in ribosome biogenesis^58^. Also, lysine residues are frequently enriched in IDRs of proteins associated with cytoplasmic protein/RNA granules and are known to promote biomolecular condensation through multivalent electrostatic interactions^59^. Consistent with this, lysine is highly abundant in MAP2/Tau family proteins^60–62^ and other MAPs^63, 64^, where it mediates interactions with NAs and MTs, and is subject to regulatory post-translation modifications, such as acetylation, that modulate MT binding and phase separation behavior^59^. Guided by this low complexity sequence grammar, we initially tested a model peptide-NA condensate system formed by a short multivalent lysine-rich repeat polypeptide [KGKGG]_5_ and a homopolymeric single-stranded DNA [dT40]. These components assemble into spherical, liquid-like condensates in vitro under physiologically relevant buffer conditions (**Figure 1B, C**).

**Figure 1.**
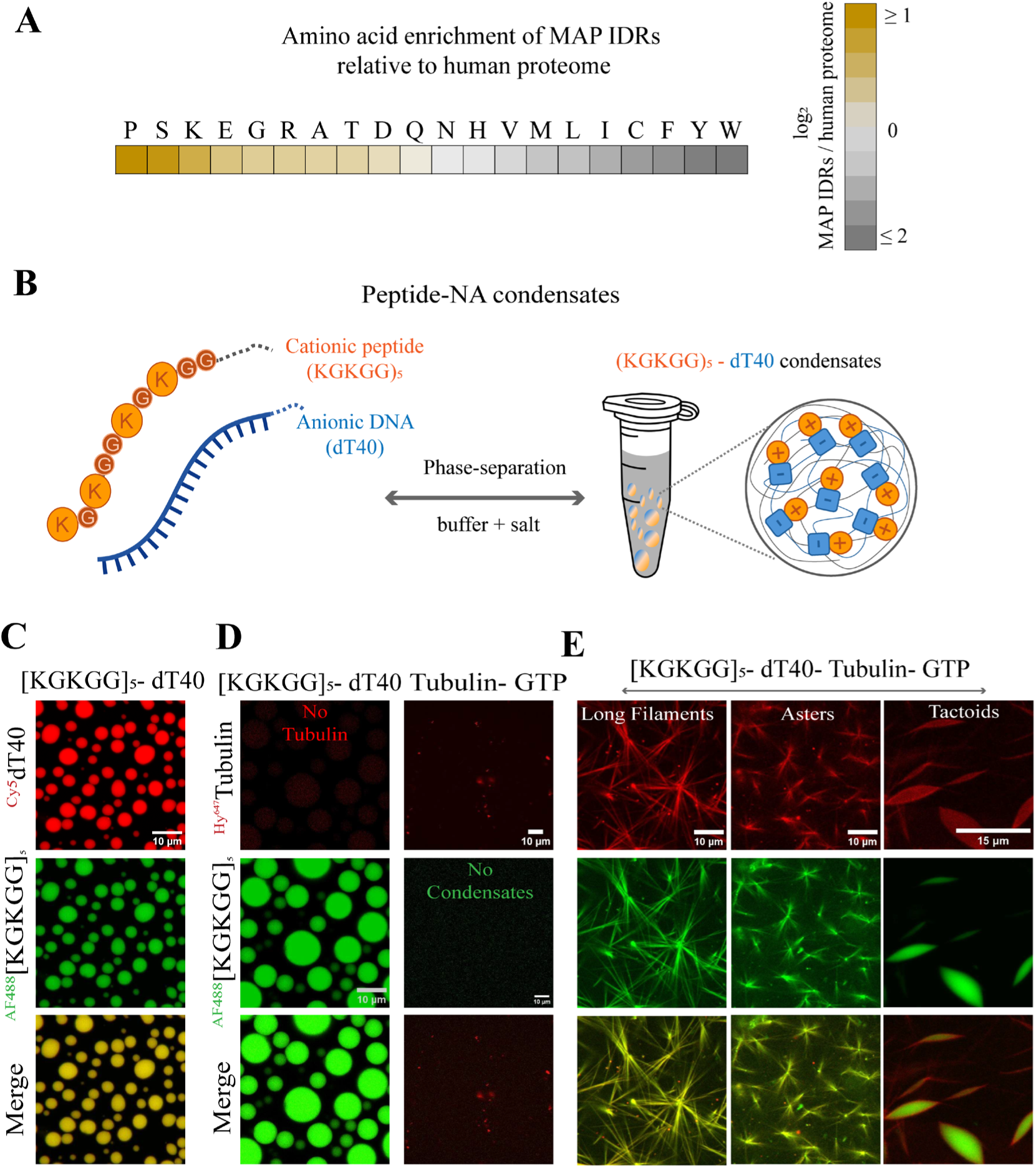
Synthetic peptide-NA condensates catalyze microtubule (MT) assembly. **(A)** Heatmap of the enrichment of indicated amino acids in human MAP IDRs relative to the human proteome. **(B)** A schematic of the model peptide-NA condensate system showing the sticker-spacer architecture of associative peptide and ssDNA chains. **(C)** Representative confocal fluorescence micrographs of [KGKGG]_5_–dT40 condensates formed at 2.5 mg/mL [KGKGG]- (green, middle) with 500 nM ^AF488^[KGKGG]_5_ and 1.25 mg/mL dT40 with 500 nM ^Cy5^dT40 (red, top) and overlay (merge, bottom). **(D)** Representative confocal fluorescence micrographs of [KGKGG]_5_–dT40 condensates formed at 2.5 mg/mL [KGKGG]_5_ (green, middle) and 1.25 mg/mL dT40 with 500 nM ^AF488^[KGKGG]_5_, without any tubulin or GTP (left panels); and 10 µM tubulin (red, top) with 900 nM ^Hy647^Tubulin and 2 mM GTP without any condensates (right panels). **(E)** Representative confocal fluorescence micrographs showing organization of MT bundles into long filaments, asters, and tactoids in the presence of [KGKGG]_5_–dT40 condensates, with 10 µM tubulin (red, top), 2 mM GTP, 2.5 mg/mL [KGKGG]_5_ (green, middle), 1.25 mg/mL dT40, and overlay (merge, bottom). Samples imaged 60 min after tubulin-GTP addition.

To assess whether [KGKGG]_5_-dT40 condensates could promote tubulin polymerization, we employed a fluorescence microscopy-based MT assembly assay^50, 53^ by adding tubulin (10 µM) together with GTP (2 mM) to pre-formed condensates. The condensates were visualized using AF488-labeled [KGKGG]_5_ (1.2 mol % of the total [KGKGG]_5_) and MTs are visualized by HiLyte 647-labeled tubulin (^Hy647^Tubulin=10% of the total tubulin). We confirmed that this tubulin concentration was much lower than the threshold required for spontaneous MT self-assembly *in vitro* under our experimental buffer conditions lacking condensates (**Figure 1D**). Remarkably, in the presence of [KGKGG]_5_-dT40 condensates, tubulin-GTP addition triggered formation of MT filaments (**Figure 1E**). Growing MTs seem to exert mechanical forces that reshape condensates into diverse morphologies, including tactoids (**Figure 1E; Video S1; Figure S1**). After an hour, dense networks of long MTs formed, including aster-like morphologies, with complete colocalization of peptide and tubulin signals, and no distinct condensates remained in solution (**Figure 1E**).

Robust MT assembly required the presence of all four components (peptide, NA, tubulin, and GTP) in this multicomponent system (**Figure S2**). Of note, minor changes in the experimental protocol, such as the order of component addition, yielded slightly variable outcomes in MT filament density and bundling (**Figure S3**); however, the primary observation that synthetic [KGKGG]_5_-dT40 condensates support robust MT assembly remained unchanged.

To more precisely track MT growth and droplet dissolution, we repeated these experiments at lower concentrations of tubulin-GTP (5 µM tubulin, 1 mM GTP) and observed an enrichment of tubulin fluorescence signal at the condensate interface, followed by MT filament growth from the interface and progressive shrinkage of the condensate (**Video S2; Figure S1B**). In contrast, the addition of equal volumes of buffer and GTP alone did not alter condensate morphology (**Video S3**). We also tested the ability of [KGKGG]_5_-NA condensates to support MT filament formation using ssDNA and RNA known to adopt stable secondary structures, in place of unstructured homopolymeric dT40, given that NA sequence and structure are important determinants of the physical properties and function of biological condensates^65, 66^. We observed robust MT formation catalyzed by these condensates (**Figure S4**). Tubulin partitioning and MT localization were observed at the condensate interface and in the surrounding dilute phase, with no visible MTs present within condensates. Collectively, these results demonstrate that Lys-rich peptide-NA condensates can robustly catalyze tubulin polymerization into MTs with complex morphologies, such as long filaments, asters, and tactoids, and that the condensate interface likely plays an important role in MT nucleation and assembly (**Video S2; Figure S1B; Figure S4**).

### Condensate mixture stoichiometry can regulate MT filament formation

Previous studies from our group and others have shown that changes in the NA-to-protein stoichiometry in the bulk can affect the condensate interfacial properties, driving a reentrant phase transition accompanied by interfacial charge inversion at the excess NA-to-peptide conditions^56, 57, 67, 68, 69^. Employing solution turbidity measurements, we confirmed that the [KGKGG]_5_-dT40 mixture exhibits reentrant phase behavior (**Figure S5)**. Using a fixed peptide concentration (2.5 mg/mL) and variable [dT40]-to-[peptide] mixing ratio, we monitored condensates via fluorescence microscopy, revealing that condensates at [dT40]:[peptide] ratio of 0.4 are significantly larger, and occupied a greater surface area on the coverslip than those prepared at the lower or higher mixing ratios (**Figure 2A**). Brightfield microscopy corroborated these results (**Figure S5**), which revealed that the condensates formed between 0.1 < [dT40]:[peptide] < 2.0.

**Figure 2.**
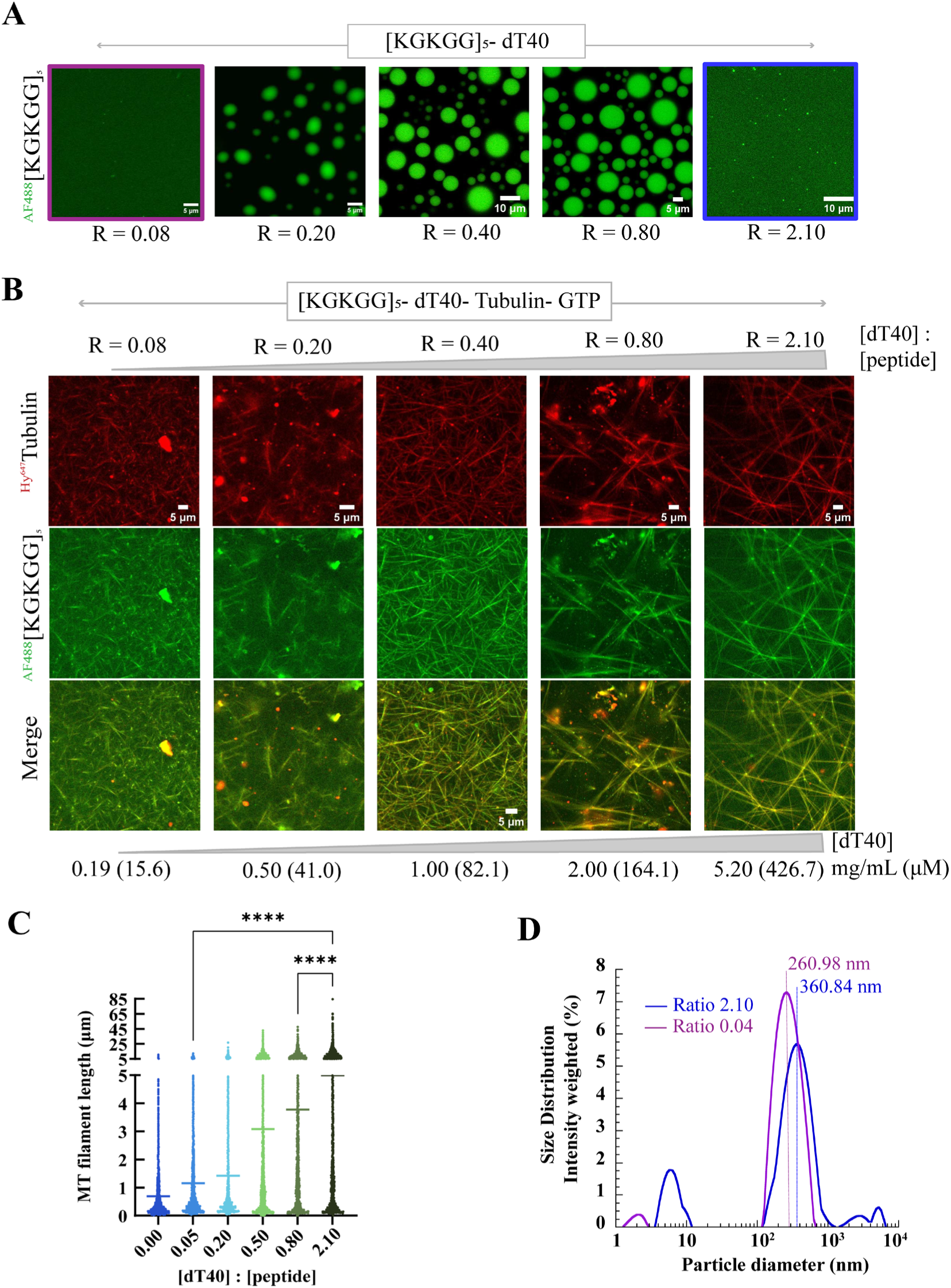
Condensate mixture stoichiometry can regulate MT filament formation. **(A)** Representative fluorescence micrographs showing reentrant phase behavior of [KGKGG]_5_–dT40 condensates as the concentration of dT40 is continuously varied between 0.19−5.2 mg/mL, whereas the concentration of [KGKGG]_5_ was fixed at 2.5 mg/mL. Panels from left to right indicate these different [dT40]-to-[peptide] mixing ratios (R). Absence of micron-scale condensates at the lowest (purple square) and highest (blue square) [dT40]:[peptide] ratio is noted. **(B)** Representative fluorescence micrographs showing variation in MT filament assembly as a function of [dT40]:[peptide] ratio. [KGKGG]_5_ was fixed at 2.5 mg/mL, while ssDNA varied between 0.19−5.2 mg/mL, with 10 µM tubulin, and 2 mM GTP. The corresponding concentrations of ssDNA in µM are indicated in parentheses. Samples imaged 60 min after tubulin-GTP addition. **(C)** MT filament length quantification plot corresponding to panel (B). Horizontal lines at each [dT40]:[peptide] ratio represent the mean values. The individual data points represent measurements based on three to four independent replicates. Sample size: [dT40]:[peptide] ratio (R) = 0.00, n = 3234; R = 0.05, n = 4804; R = 0.20, n = 4777; R = 0.50, n = 4214; R = 0.80, n = 3056, and R = 2.10, n = 4665. Statistical significance was determined using a one-way ANOVA Dunn’s multiple comparisons test between the individual ratios (* means p < 0.05, ** means p < 0.01, *** means p < 0.001, **** means p < 0.0001). The associated *P* values are shown only for R = 0.05 vs. R = 2.1 and R = 0.80 vs. R = 2.10. **(D)** Size distribution of [KGKGG]_5_–dT40 clusters at the lowest (purple) and highest (blue) [dT40]:[peptide] ratio obtained from DLS measurements from two independent samples.

To test whether condensate-mediated MT assembly is impacted by the bulk mixture composition across the reentrant phase boundary, we examined MT assembly in [KGKGG]_5_-dT40 condensates prepared across a range of nucleotide-to-peptide ratios ([dT40]:[peptide]) from 0.08 to 2.1. Surprisingly, we found that condensates formed at the excess peptide conditions ([dT40]:[peptide] = 0.08) catalyzed the nucleation of distinctively short MT seeds, whereas condensates with excess dT40 ([dT40]:[peptide] = 2.1) catalyzed the formation of long branched MT filaments under otherwise identical conditions (**Figures 2B, C; Figures S6-7**).

To determine the size of peptide-NA complexes at the lowest and highest mixing ratios, we performed dynamic light scattering (DLS) measurements. DLS revealed the presence of submicron-sized complexes (mean diameters ∼250–380 nm) under these conditions (**Figure 2D**), suggesting the existence of nanoscale peptide–NA clusters. Strikingly, time-lapse fluorescence imaging of MT assembly revealed that these nanoscale clusters, formed at [dT40]:[peptide] = 2.1, rapidly nucleated MTs, with long filaments emerging within 15 min of tubulin-GTP addition (**Video S4; Figure S6**). Together, these results demonstrate that the stoichiometry of the condensate mixture governs MT assembly efficiency and the morphology of the resultant MT filaments, likely by modulating condensate interfacial properties. In particular, nanoscale peptide–NA clusters formed under NA-rich conditions, which carry excess NA on their interface and hence have negatively charged interface (**Figure S5B**), act as highly active sites for long MT filament formation.

### Condensates formed by peptides with distinct sequence grammars differentially regulate tubulin mobility and MT growth

Having established that condensates formed by Lys-rich cationic peptide and ssDNA complexes can act as hubs for MT assembly, we next investigated how the chemical grammar of the cationic peptide and condensate microenvironment, such as the viscoelasticity of the fluid network, influences MT nucleation and growth. Amino acid enrichment analysis of MAP IDRs (**Figure 1A**) indicates a modest preference for Lys over Arg. Although Lys and Arg are both singly positively charged at physiological pH and act as sticker residues in NA-associated phase separation, Arg generally binds NAs more strongly than Lys^21, 56, 70, 71^. In addition to the differential interactions encoded by cationic “stickers” (R vs. K), the intervening spacer residues (G vs. P) can further modulate NA binding and the network strengths of peptide-NA condensates^21, 22, 57^. These collective differences in molecular-level interactions provide a key handle to develop synthetic peptide-NA condensates with programmable viscoelasticity through modular peptide design^21, 22^.

To examine whether and how the material properties imparted by the fluid network of these condensates regulate MT assembly, we employed three distinct condensate systems formed by either (KGKGG)_5_, (RGRGG)_5_, or (RPRPP)_5_ polypeptides and dT40. We first quantified the material properties of these two-component peptide–NA condensates using video particle tracking (VPT) nanorheology^7, 21^. The ensemble-averaged mean squared displacement (MSD) profiles obtained from VPT measurements using 200-nm fluorescently labeled probe particles revealed distinct differences in the material properties and exhibited trends like those previously observed for two-component peptide–RNA systems^21, 22^. Specifically, the viscosity of condensates formed with RGRGG-repeat peptides was ∼150-fold higher than the condensates formed by KGKGG-repeat peptide (viscosities, η ∼15.04 ± 1.13 Pa.s vs. 0.1 ± 0.01 Pa.s; **Figure 3A, B**), suggesting a dominantly viscous microenvironment within the (KGKGG)_5_-dT40 condensates. We further characterized the average mesh size of the viscoelastic networks in these condensates using a variable-size dextran recruitment assay^21, 72^, revealing a larger mesh size for (KGKGG)_5_-dT40 condensates compared to the (RGRGG)_5_-dT40 condensates (**Figure S8**). The (RPRPP)_5_-dT40 condensates displayed an intermediate viscosity (η ∼1.93 ± 0.24 Pa.s), which is ∼20-fold higher than (KGKGG)_5_-dT40 condensates and ∼8-fold lower than (RGRGG)_5_-dT40 condensates (**Figure 3A**, **B**).

**Figure 3.**
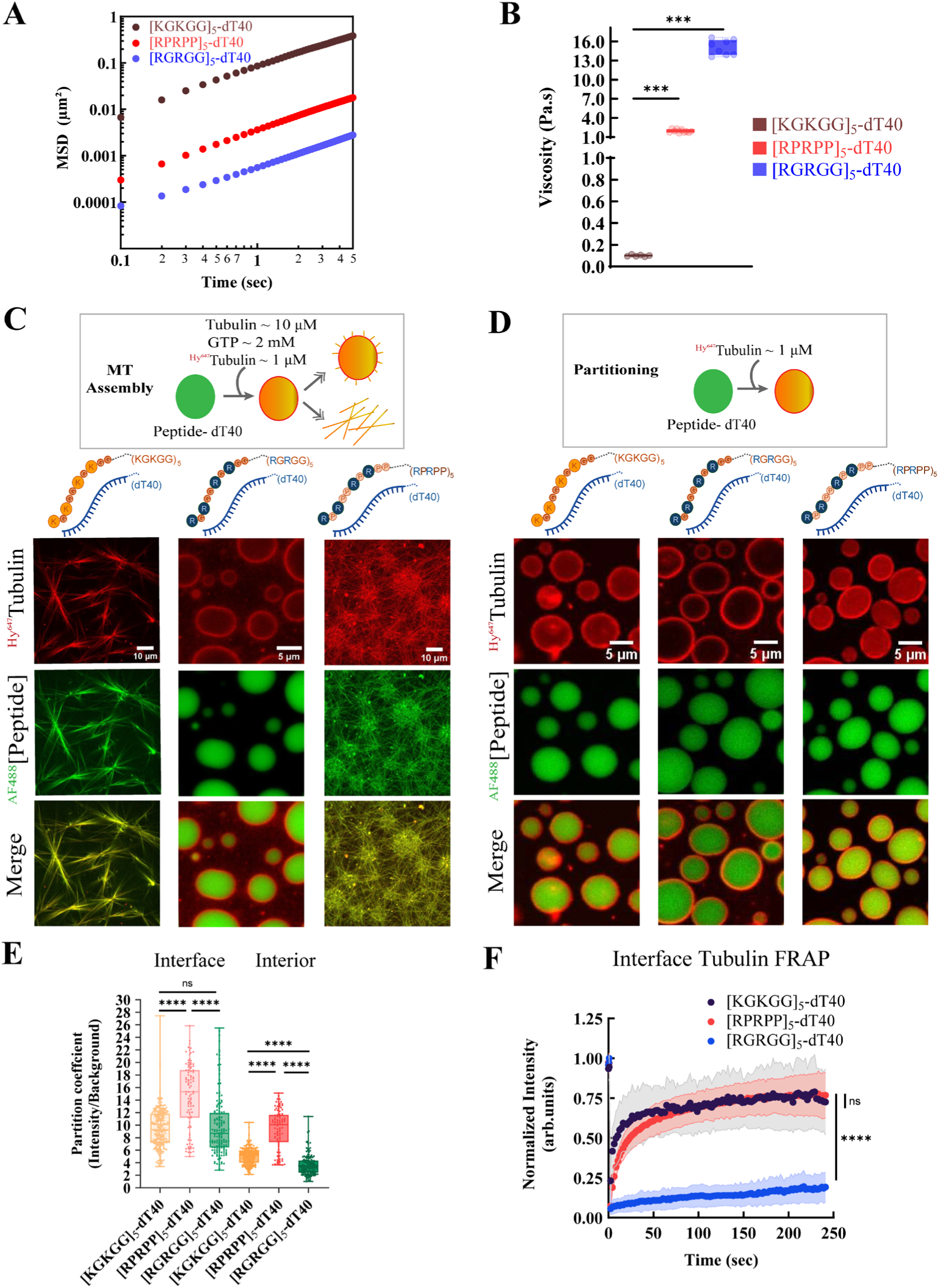
Condensates formed by peptides with distinct sequence grammars differentially regulate tubulin mobility and MT growth. **(A)** The ensemble-averaged mean square displacement (MSD) estimated from passive motion of 200 nm polystyrene beads within [KGKGG]_5_-dT40 condensates (brown), [RGRGG]_5_-dT40 condensates (blue), and [RPRPP]_5_-dT40 (red), using video particle tracking (VPT) nanorheology (see Methods section for further details). Shifts in MSDs (top to bottom) signify decreasing diffusivity of beads inside the condensates. **(B)** Variation in terminal viscosity values of [KGKGG]_5_-dT40 (brown), [RGRGG]_5_-dT40 (blue), and [RPRPP]_5_-dT40 (red) condensates estimated from fitting longer lag time ensemble-averaged MSDs to 𝑀𝑆𝐷(𝜏) = 4𝐷𝜏^𝛼^ + 𝑁 . [KGKGG]_5_-dT40, [RGRGG]_5_-dT40, and [RPRPP]_5_-dT40 condensates show a viscosity of 0.1 ± 0.01 Pa.s, 15.04 ± 1.13 Pa.s, and 1.93 ± 0.24 Pa.s, respectively. The data is shown for n = 12−15 condensates across 3 independent sample replicates. **(C)** The schematic on top depicts the MT assembly assay with GTP. Representative confocal fluorescence micrographs showing organization of MT bundles into long filaments in the presence of [KGKGG]_5_–dT40 condensates (left panels); short MT asters form in the presence of [RGRGG]_5_–dT40 condensates (middle panels); and the organization of long MT filaments in the presence of [RPRPP]_5_–dT40 condensates (right panels); The condensate composition and sample conditions are as reported in Fig 1D. **(D)** The schematic on top depicts the tubulin partitioning experiments without GTP. Fluorescence microscopy images showing the recruitment behavior of tubulin into [KGKGG]_5_-dT40 (left panels), [RGRGG]_5_-dT40 condensates (middle panels), and [RPRPP]_5_-dT40 condensates (right panels). The condensate composition and sample conditions are as in Fig. 3C, but samples contained only 900 nM ^Hy647^Tubulin (no excess unlabeled tubulin). **(E)** Tubulin partition coefficient measurements at the interface and interior of [KGKGG]_5_-dT40 (Interface: light orange, Interior: dark orange), [RGRGG]_5_-dT40 condensates (Interface: light green, Interior: dark green), and [RPRPP]_5_-dT40 (Interface: light pink, Interior: dark pink). The individual data points represent measurements based on two to three independent replicates. Sample size: [KGKGG]_5_-dT40 (Interface & Interior: n = 245), [RGRGG]_5_-dT40 (Interface & Interior: n = 161), and [RPRPP]_5_-dT40 (Interface & Interior: n = 96). For the interfacial partitioning, the associated *P* values are p = 0.3011 for [KGKGG]_5_-dT40 vs. [RGRGG]_5_-dT40 condensates; p < 0.0001 for all other pairwise combinations. **(F)** FRAP intensity time-traces of tubulin probe (^Hy647^Tubulin) measured at the interface of condensates formed by dT40 ssDNA and [KGKGG]_5_ (brown), [RGRGG]_5_ (blue), and [RPRPP]_5_ (pink), respectively. Sample conditions as described in (E). Solid circles are the average of 5-7 FRAP curves for each peptide-DNA combination across 2 independent sample replicates. Error bars in the curves represent one standard deviation (±1 s.d.). Statistical significance was determined using a one-way ANOVA Dunn’s multiple comparisons test between the individual conditions. In B and E, the box represents the interquartile range from the 25^th^ to the 75^th^ percentile, with the middle line showing the median, the whiskers showing the minimum to maximum values, and the individual datapoints plotted as circular symbols. Statistical significance was determined using an unpaired two-tailed Student’s t-test (B) and a one-way ANOVA Dunn’s multiple comparisons test (E) between the samples (“ns” means non-significant, * means p < 0.05, ** means p < 0.01, *** means p < 0.001, **** means p < 0.0001).

We next assessed how these condensates, formed by peptides with distinct sequence grammar and material properties, influence MT polymerization. Under identical reaction conditions, the highly viscoelastic (RGRGG)_5_-dT40 condensates failed to assemble long MT filaments (**Figure 3C**). Time-lapse fluorescence imaging revealed that, unlike (KGKGG)_5_-dT40 condensates, the (RGRGG)_5_-dT40 condensates remained stable (**Video S5**). Interestingly, the smaller (RGRGG)_5_-dT40 condensates exhibited shorter MT filament projections than larger condensates, but filament growth was arrested at this stage, with no further increase in MT filament length observed over time (**Figure S9A**). This remarkable difference in MT polymerization between (KGKGG)_5_-dT40 and (RGRGG)_5_-dT40 condensates indicates either a profound role of condensate material properties in MT polymerization and/or that lysine-rich, but not arginine-rich, peptides are selectively capable of supporting MT assembly. To disentangle these two possibilities, we tested the (RPRPP)_5_-dT40 condensates, which retain the same number of arginine but have different spacers and significantly lower viscosity than (RGRGG)_5_-dT40 condensates. We observed that, like (KGKGG)_5_-dT40 condensates, (RPRPP)_5_-dT40 condensates can also catalyze MT assembly with long filaments forming under identical experimental conditions (**Figure 3C**). These results suggest that the material properties of host condensates, encoded by the peptide sequence grammar, play a regulatory role in MT polymerization.

To probe the mechanistic basis for the observed variation in MT assembly, we examined tubulin partitioning in these three peptide-NA condensate systems. Using fluorescence microscopy, we determined the partition coefficients of labeled tubulin at the condensate interface as well as the interior (**Figure 3D, E**). The partition coefficient is calculated as the ratio of the fluorescence intensity of tubulin either at the interface (k_interface_) or in the interior (k_interior_) to that in the dilute phase. Partition coefficients report on the transfer free energy of a macromolecule from the dilute phase to the dense phase, which is determined, in part, by intermolecular interactions between the client and condensate-forming scaffolds^73^. Importantly, in these three peptide-NA condensate systems, fluorescence intensity of labeled tubulin was observed to be selectively enriched at the condensate interface relative to the condensate interior. Although interfacial tubulin enrichment was comparable between (KGKGG)_5_-dT40 (k_interface_ = 9.45 ± 3.15; η = 0.1 ± 0.01 Pa.s) and (RGRGG)_5_–dT40 condensates (k_interface_ = 9.78 ± 4.75; η = 15.04 ± 1.13 Pa.s), enrichment was higher in (RPRPP)_5_–dT40 condensates (k_interface_ = 14.72 ± 5.21; η = 1.93 ± 0.24 Pa.s). In contrast, tubulin partitioning into the condensate interior was greater for (KGKGG)_5_-dT40 (k_interior_ = 5.07 ± 1.31) and (RPRPP)_5_-dT40 condensates (k_interior_ = 9.36 ± 3.19) than for (RGRGG)_5_–dT40 condensates (k_interior_ = 3.59 ± 1.62). This differential partitioning of tubulin likely arises from differences in competitive interactions between tubulin and the repeat peptides for ssDNA binding and/or the differences in condensate mesh size (**Figures S8, S10**).

The interface of biomolecular condensates has recently been reported to promote nucleation of protein fibrils^74–76^. Since tubulin shows selective partitioning to the condensate interface in our condensate systems, the interface can serve as the site for MT nucleation. Therefore, we wondered whether the observed alterations in MT filament formation across different condensate systems arise from altered tubulin diffusion dynamics at the interface. Unlike nanorheology, which quantifies condensate viscoelasticity through the Brownian motion of embedded 200-nm beads, fluorescence recovery after photobleaching (FRAP) measures the diffusion dynamics of fluorescently labeled proteins within a relatively larger bleached region (> 400 nm). Consequently, FRAP reflects the combined effects of molecular mobility, intermolecular interactions, and network viscoelasticity^22^. FRAP measurements revealed that the tubulin mobility was significantly higher at the interface of (KGKGG)_5_-dT40 condensates (τ_1/2_ = 10.38 ± 1.22 sec; mobile fraction = 71.20 ± 26.10%) when compared to (RGRGG)_5_-dT40 condensates (τ_1/2_ = 444.26 ± 88.44 sec; mobile fraction = 13.71 ± 7.18%) (**Figure 3F; Figure S11; Tables S1-2; Videos S6, S8**). Remarkably, tubulin mobility at the interface of (RPRPP)_5_-dT40 condensates (τ_1/2_ = 17.50 ± 0.63 sec; mobile fraction = 75.20 ± 15.5%) closely matches that observed in (KGKGG)_5_-dT40 condensates (**Figure 3F; Figure S11; Tables S1-2; Videos S6, S10**). Similar trends in tubulin mobility were also observed for FRAP measurements performed in the interior of these three peptide-NA condensate systems (**Figure S12D; Videos S7, S9, S11**). These FRAP results, therefore, suggest a direct correlation between interfacial tubulin mobility and MT assembly aided by synthetic peptide-NA condensates. Collectively, these results suggest that the material properties of peptide-NA condensates encoded by the peptide sequence grammar can differentially regulate tubulin partitioning at the interface, diffusivity, and ultimately MT assembly.

### Condensate material state regulates MT assembly

Tuning the condensate’s material properties by varying the peptide sequence grammar (Lys vs. Arg; **Figure 3**) can alter peptide−tubulin interactions (**Figure S10**). To decouple such effects from the role of condensate material properties, we adopted an orthogonal approach to tune condensate viscoelasticity without modifying the peptide sequence grammar. We previously showed that increasing the length of the condensate-scaffolding NA, while keeping the identity of the scaffolding peptide constant, provides an independent route to controlling condensate viscoelasticity^22^. Following this strategy, we varied the poly-dT chain length from 20 to 200 nucleobases and combined these with the (KGKGG)_5_-peptide. The resulting condensate systems span an ∼8-fold range in viscosity (η = 0.1 ± 0.01 Pa.s to 0.73 ± 0.06 Pa.s; **Figure 4A**, **B**). Additionally, to isolate the contribution of peptide sequence identity from viscosity, we also substituted 50% of lysine residues in (KGKGG)_5_ peptide with tyrosine, yielding (KGYGG)_5_-dT40 condensates that retained partial sequence grammar to (KGKGG)_5_-dT40 condensates but showed a ∼ 70-fold increase in viscosity (η = 6.95 ± 1.53 Pa.s; **Figure 4A**, **B**).

**Figure 4.**
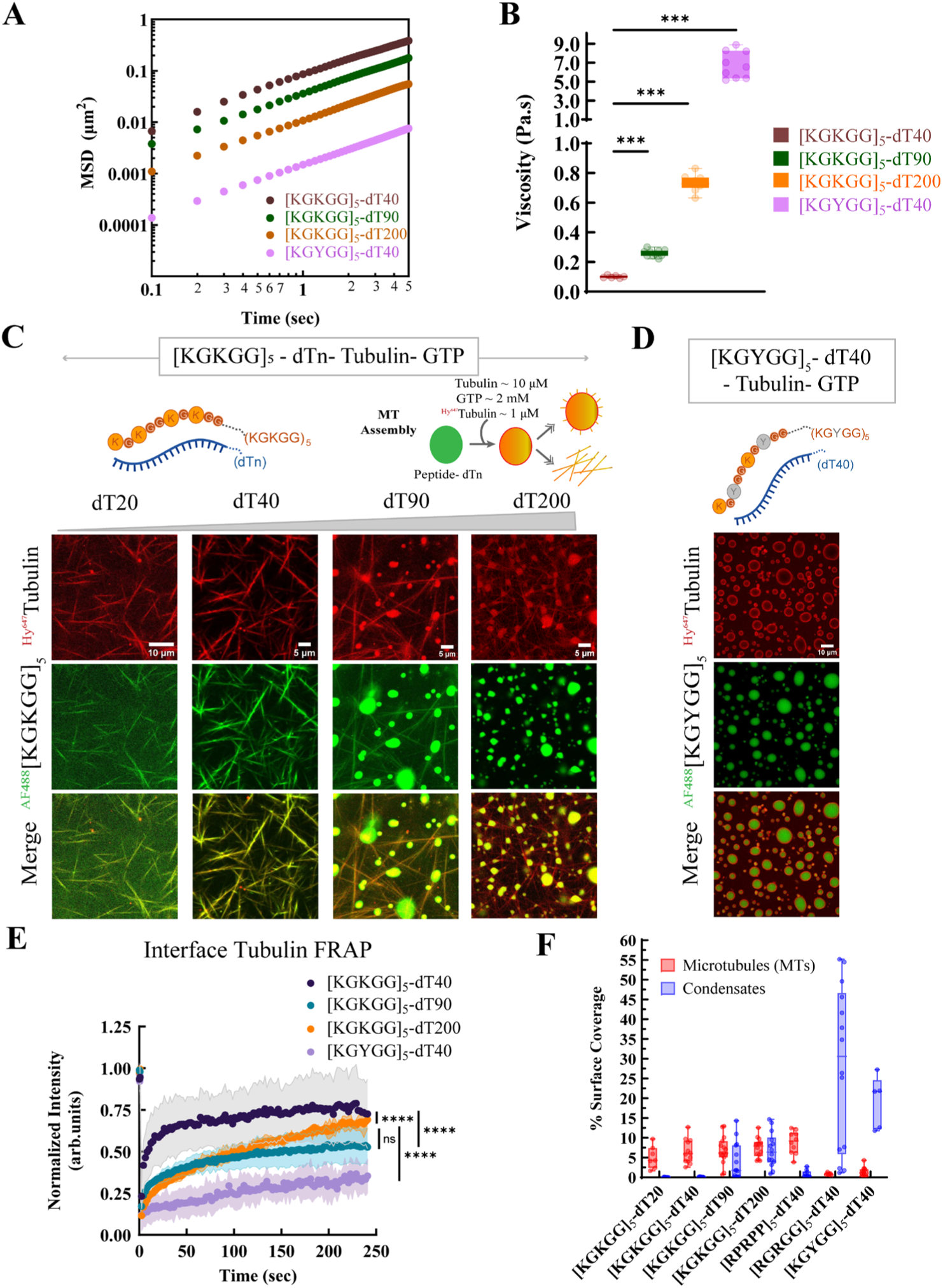
Condensate material state regulates MT assembly. **(A)** The ensemble-averaged mean square displacement (MSD) estimated from the passive motion of 200 nm polystyrene beads within [KGKGG]_5_-dT40 condensates (brown), [KGKGG]_5_-dT90 condensates (green), [KGKGG]_5_-dT200 (orange), and [KGYGG]_5_-dT40 (purple) using VPT nanorheology. Shifts in MSDs (top to bottom) signify decreasing diffusivity of beads inside condensate. **(B)** Variation in terminal viscosity values across the different peptide-NA condensate systems is shown. [KGKGG]_5_-dT40, [KGKGG]_5_-dT90, [KGKGG]_5_-dT200, and [KGYGG]5-dT40 condensates show viscosities of 0.1 ± 0.01 Pa.s, 0.26 ± 0.02 Pa.s, 0.73 ± 0.06 Pa.s, and 6.95 ± 1.53 Pa.s, respectively. **(C)** A schematic and representative confocal fluorescence micrographs showing the variation in organization of MT filaments in the presence of different condensate systems with distinct viscosities, arranged in order of increasing condensate viscosity from left to right. The condensate composition and sample conditions are as reported in Figs. 1D & 3C. Samples were imaged 90 min after tubulin-GTP addition. **(D)** Representative confocal fluorescence micrographs showing the lack of MT filaments in the presence of [KGYGG]_5_-dT40 condensates. Reaction conditions as described in (C). **(E)** FRAP intensity-time traces of tubulin probe (^Hy647^Tubulin) measured at the interface of condensates formed by [KGKGG]_5_-dT40 (brown), [KGKGG]_5_-dT90 (green), and [KGKGG]_5_-dT200 (orange), and [KGYGG]_5_-dT40 (purple). Sample conditions as described in (Fig. 3E, F). Solid circles are the average of 4-5 FRAP curves for each peptide-DNA combination across two independent sample replicates. Error bars in the curves represent one standard deviation (±1 s.d.). Statistical significance was determined using a one-way ANOVA Dunn’s multiple comparisons test between the individual conditions, where ‘ns’ means non-significant, **** means P ≤ 0.0001. **(F)** Plot showing percentage (%) surface area coverage of MT filaments (red), and condensates (blue) corresponding to panels (C, D; Fig 3C). The center line represents the mean. The individual data points based on measurements from at least three independent replicates for each condition are shown. Sample size: [KGKGG]_5_-dT20, 7 frames; [KGKGG]_5_-dT40, 12 frames; [KGKGG]_5_-dT90, 15 frames; [KGKGG]_5_-dT200, 14 frames; [RPRPP]_5_-dT40, 10 frames; [RGRGG]_5_-dT40, 14 frames; [KGYGG]_5_-dT40, 5 frames. Statistical significance was determined using a one-way ANOVA Dunn’s multiple comparisons test between the individual conditions. MT surface coverage showed: “ns” (p > 0.999) among [KGKGG]_5_-dT20 / dT40 / dT90 / dT200 samples; “**” (p < 0.01) between [KGKGG]_5_-dT40/dT90 vs. [KGYGG]_5_-dT40 samples; “***” (p < 0.001) between [KGKGG]_5_-dT200 or [RPRPP]_5_-dT40 vs [KGYGG]_5_-dT40 samples; “*” (p < 0.05) between [KGKGG]_5_-dT40 / dT90 vs [RGRGG]_5_-dT40 samples; “**” (p < 0.01) between [KGKGG]_5_-dT200 or [RPRPP]_5_-dT40 vs [RGRGG]_5_-dT40 samples; Condensate surface coverage showed: “ns” (p > 0.999) among [KGKGG]_5_-dT20 / dT40 vs [KGKGG]_5_-dT90 samples; “***” (p < 0.001) between [KGKGG]_5_-dT20 / dT40 vs [KGKGG]_5_-dT200 samples or [KGYGG]_5_-dT40 samples; “****” (p < 0.0001) between [KGKGG]_5_-dT20 / T40 vs [RGRGG]_5_-dT40 samples; “**” (p < 0.01) between [RPRPP]_5_-dT40 vs [RGRGG]_5_-dT40 samples.

We next assessed MT polymerization in the presence of these condensates with variable material and physical properties. Under identical reaction conditions, we observed a modest decline in MT assembly efficiency with increasing ssDNA chain length. Although (KGKGG)_5_-dT200 condensates supported MT polymerization, thick filaments were absent, and more condensate-like assemblies were observed (**Figure 4C, F**). Notably, (KGYGG)_5_-dT40 condensates failed to assemble MT filaments (**Figure 4D, F**).

Interestingly, tubulin partitioning across these condensate systems did not entirely mirror the observed trends in condensate-mediated MT nucleation and growth. For instance, although MT assembly was more efficient in (KGKGG)_5_-dT200 condensates than in (RGRGG)_5_-dT40 condensates, interfacial tubulin enrichment was at least ∼3-fold lower in the former. (k_interface_ = 3.02 ± 1.43 vs 9.78 ± 4.75; **Figure 4D; Figure S12A-C**). However, FRAP measurements revealed that the observed outcome of MT filament formation correlates well with the degree of tubulin mobility within and at the interface of these peptide-NA condensates (**Figure 4E; Figure S11, S12D; Tables S1-2; Videos S12-15**). We note that due to the lack of tubulin enrichment within the interior of (KGYGG)_5_-dT40 condensates, and the absence of distinct interfacial accumulation, our FRAP analysis was confined to the interfacial region where tubulin mobility could be most reliably assessed, and it was observed to be significantly lower as compared to the (KGKGG)_5_-dT40 condensates (τ_1/2_ = 247.9 ± 44.5 sec vs. 10.38 ± 1.22 sec; mobile fraction: 27.5 ± 12.8% vs. 71.2 ± 26.1%, respectively; **Figure 4E; Figure S11; Tables S1-2; Video S16**).

Together, these results suggest that condensate-mediated regulation of MT assembly depends on a complex interplay between peptide sequence grammar and condensate material state. While higher condensate viscoelasticity broadly suppresses MT filament formation, peptide sequence–encoded interactions further fine-tune the balance between condensate stability, tubulin dynamics, and MT assembly.

### A minimal coarse-grained stochastic kinetics model provides mechanistic insight into the condensate-interface directed MT assembly dynamics

To interpret how condensate material properties encode distinct MT assembly outcomes, we developed a coarse-grained stochastic model of MT nucleation and growth at a spherical condensate surface (**Figure S13**; full description in Methods Section; Supplementary Note S1; **Figure 5A**). While classical stochastic MT models^77, 78^ treat monomer supply solely through dilute phase diffusion, our model introduces two key extensions: a two-dimensional tubulin field on the condensate surface, whose lateral mobility is controlled by interfacial viscosity, and a Kramers-type detachment activation barrier^79^ that links condensate material properties to MT escape kinetics. Tubulin is represented as a surface density field 𝑛(𝐮, 𝑡) that is replenished from the dilute phase by adsorption and redistributed laterally by surface diffusion. MT nucleation, elongation, and tip detachment are simulated using stochastic Gillespie kinetics^80^ with rates coupled to the local surface density. The effective condensate interface viscosity 𝜂_eff_ is the single parameter encoding sequence-encoded property of the condensate interface. 𝜂_eff_simultaneously sets the effective 2D surface diffusivity of tubulins via 𝐷_surf_ = 1/𝜂_eff_(lateral tubulin mobility) and the activation barrier 𝑒^−𝛼𝜂e𝐹𝐹^ controlling the dissociation rate of the MT tip^79^ from the condensate interface, where α is a phenomenological factor that converts interfacial viscosity to adhesion free energy. High 𝜂_eff_ therefore corresponds to dense, cohesive interfaces characteristic of RGRGG-type sequences, while low 𝜂_eff_ represents more liquid-like, weakly adhesive interfaces characteristic of KGKGG-type sequences.

**Figure 5:**
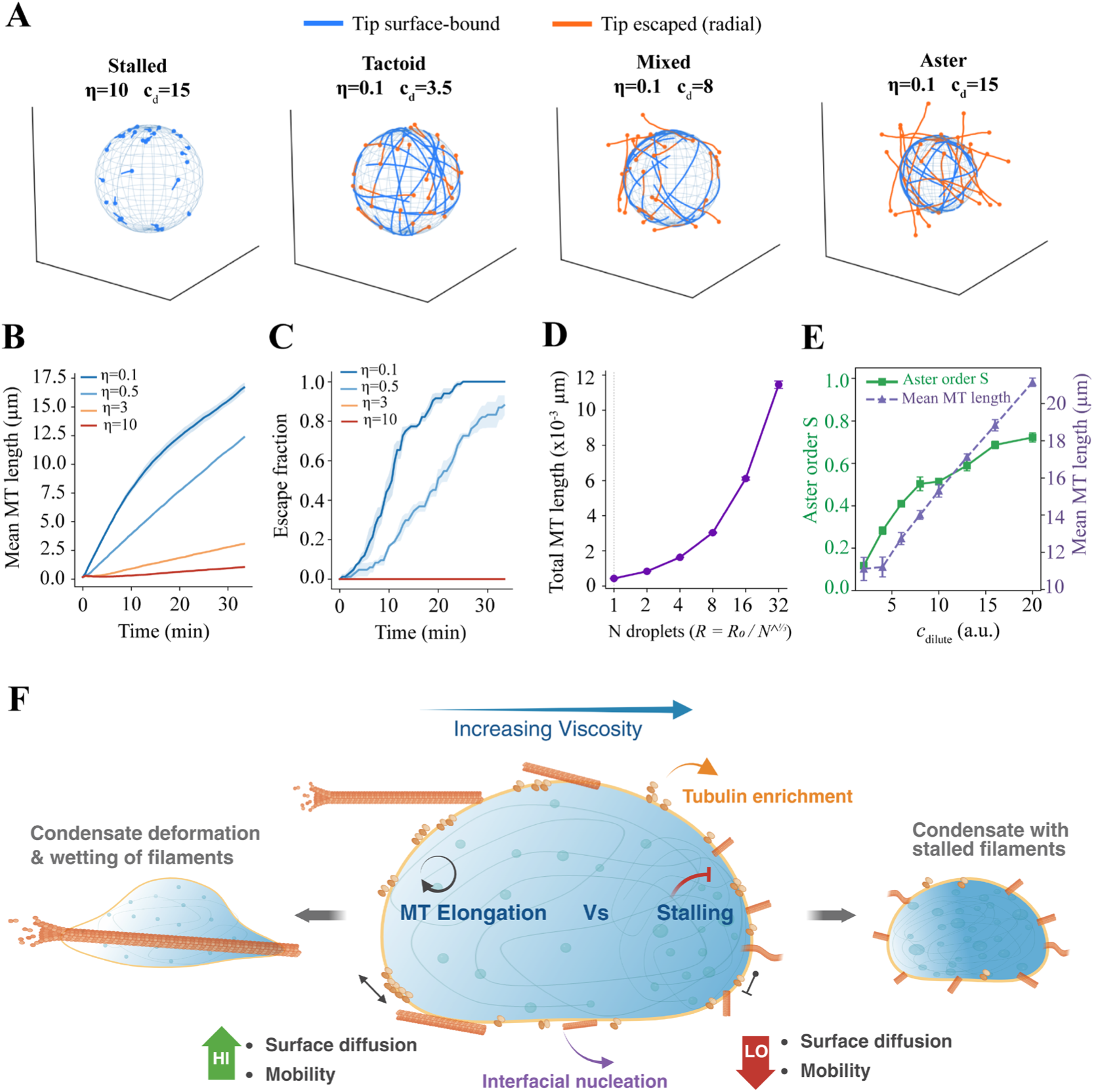
Condensate interfacial rheology and tubulin partitioning jointly control MT assembly morphology and kinetics. **(A)** Simulation snapshots at 𝑡 = 2000 s illustrating four assembly regimes. Blue filaments have tips in contact with the condensate surface; orange filaments have escaped and grow radially into the bulk. From *left* to *right*: Stalled (high 𝜂_eff_, high 𝑐_dilute_) short, surface-arrested filaments with no escape, representative of highly viscoelastic RGRGG-type interfaces (**Video S17**); Tactoid (low 𝜂_eff_, low 𝑐_dilute_) tangentially ordered surface filaments; Mixed (low 𝜂_eff_, intermediate 𝑐_dilute_) coexistence of surface-bound and escaped filaments; Aster (low 𝜂_eff_, high 𝑐_dilute_ ) radially organized, predominantly free MTs characteristic of fluid KGKGG-type interfaces (**Video S18**). **(B)** Mean MT length over time across four interface viscosities at fixed 𝑐_dilute_ (𝑛 = 3 replicates, shading represents ± SEM). High 𝜂_eff_ stalls elongation; low 𝜂_eff_ supports rapid growth. **(C)** Surface escape fraction (1 − 𝑓_bound_) over time for the same conditions. **(D)** Total MT polymer length across 𝑁 condensate droplets at fixed total condensate volume (𝑛 = 3, error bars represent ± SEM). **(E)** Aster order parameter 𝑆 (green) and mean MT length (purple, dashed) versus dilute phase tubulin concentration at low 𝜂_eff_ (𝑛 = 3, error bars represent ± SEM). **(F)** Schematic (created with BioRender.com) depiction of the proposed model summarizing the key findings of this study: (i) Preferential partitioning of tubulin to interface of condensates; (ii) MT nucleation at the interface; (iii) regulation of MT filament growth by interfacial rheology and tubulin mobility, accompanied by (iv) deformation of less viscoelastic condensates and MT filament surface wetting by condensates or persistence of intact highly viscoelastic condensates with short, arrested filaments.

### Interface viscosity governs MT growth kinetics and surface escape

Simulations across a four-decade range of 𝜂_eff_at fixed bulk tubulin concentration (𝑐_dilute_ = 10 a.u.) reveal a clear viscosity-dependent divergence in MT assembly kinetics (**Figure 5B, C**). A surface-bound tip consumes local tubulin as it grows, creating a depletion zone. Continued elongation therefore depends on whether lateral surface diffusion can replenish the tip faster than it consumes subunits. The rate coefficient of growth 𝑘_growth_ = 𝑘_+_ 𝐷_surf_/(𝐷_surf_ + 𝐷_0_) captures this competition, where 𝑘_+_ is the intrinsic MT growth rate and 𝐷_0_ is the crossover diffusivity below which surface transport limits growth. When surface diffusion is fast (𝐷_surf_ ≫ 𝐷_0_), tubulin supply keeps pace and growth proceeds; when diffusion is slow (𝐷_surf_ ≪ 𝐷_0_), the tip becomes locally depleted of tubulin and elongation stalls.

Looking at simulation results, we see clearly that at low interface viscosity (𝜂_eff_ = 0.1), surface-bound tips grow rapidly, the diffusive-saturation factor (𝐷_surf_/(𝐷_surf_ + 𝐷_0_) approaches unity and long MTs accumulate before peeling from the surface and escaping radially into the dilute phase, producing a high escape fraction (**Figure 5C**). At high viscosity (𝜂_eff_ = 10), the same factor is suppressed by two orders of magnitude, stalling surface growth, while the activation barrier for MT dissociation simultaneously tethers tips to the interface and prevents detachment. The result is a population of short, surface-arrested filaments that neither grow nor escape, reminiscent of the kinetically stalled state observed in experiments with (RGRGG)_5_-dT40 condensates (**Figure S9B**). Overall, these two regimes are consistent with the experimentally observed contrast between (KGKGG)_5_-dT40, which supports vigorous MT nucleation and elongation, and more viscoelastic (RGRGG)_5_-dT40 condensates, where filament dynamics are markedly reduced. The intermediate viscosity conditions (𝜂_eff_ = 0.5 − 3) produce mixed morphologies with moderate growth rates and partial escape, reminiscent of (RPRPP)_5_-dT40 and (KGKGG)_5_-dT200 condensates, underscoring that the peptide sequence-based tuning of condensate material and interfacial properties regulates MT assembly dynamics continuously rather than through a binary switch.

### Condensate fragmentation non-monotonically regulates MT assembly

A key prediction of the transport-limited growth mechanism is that droplet size and number impacts the MT assembly outcome. At a fixed total condensate volume, fragmenting a single droplet into 𝑁 smaller ones of radius 𝑅 = 𝑅_0_/𝑁^1/3^ modulates the effective adsorption flux 𝑘_ads_ α 1/𝑅 and nucleation geometry. Our simulations show that total MT polymer length (summed across all 𝑁 droplets) increases sharply as 𝑁 rises from 1 to ∼8 and then saturates or slightly decreases at large 𝑁 (**Figure 5D**). Smaller droplets present a larger surface-to-volume ratio, enhancing per-droplet tubulin recruitment and nucleation, but at very large 𝑁 each droplet is too small to sustain long filaments before they geometrically outgrow the surface. Overall, these results provide a mechanistic basis for our experimental observations that (i) nanoscale peptide-NA clusters rapidly nucleated MTs and catalyzed the formation of long filaments (**Figure 2B**), and (ii) the smaller (RGRGG)_5_-dT40 condensates exhibited comparatively higher numbers of short MT filament projections than larger condensates (**Figure S9**).

### Tubulin concentration drives a tactoid-to-aster morphology transition

We next asked how tuning interfacial enrichment regulates the condensate-catalyzed MT filament morphology. At low interface viscosity, simulation reveals that increasing tubulin concentration drives a sharp transition in MT organization from tangentially ordered tactoids to radial asters (**Figure 5A, E**). At low 𝑐_dilute_, the sparse surface field supports few nucleation events, and MTs grow tangentially along the droplet surface, producing a low aster order parameter 𝑆 ≈ 0. As 𝑐_dilute_ increases, both the nucleation rate and the surface replenishment flux grow, enabling rapid filament elongation and eventual escape; MT tips detach and grow radially outward, raising 𝑆 toward 1 and increasing the mean MT length simultaneously (**Figure 5E**). The co-occurrence of increasing length and the aster order parameter mirrors the experimental observation that enhanced tubulin self-assembly produces long filaments with progressively more pronounced radial organization, and that condensate-directed polymerization can switch between bundled tactoid-like and star-like aster morphologies depending on tubulin availability at the condensate interface.

## Discussion

Using a synthetic peptide–NA system with programmable sequence grammar that controls condensate viscoelasticity, we show that condensate mechanical properties can differentially regulate tubulin dynamics and catalyze MT assembly through combined effects of material state and molecular interactions. These results suggest that condensates can act as tunable hubs whose physical and material properties directly influence MT assembly dynamics and the filament morphology. Lysine-rich, less viscoelastic condensates promote robust MT nucleation and filament elongation and bundling, whereas more viscoelastic condensates formed by arginine- or tyrosine-containing peptides restrict filament growth (**Figures 3 & 4**). This experimentally observed inverse correlation between viscoelasticity and MT assembly suggests that the condensate material state governs polymerization by modulating molecular accessibility and reaction turnover, particularly through its control over tubulin mobility. Such tunability suggests a biophysical strategy by which natural condensates may regulate cytoskeletal assembly and organization by switching between reaction-permissive and reaction-restricted regimes in response to compositional or environmental changes.

A key component of condensate-aided MT assembly is the role of the condensate interface. Tubulin preferentially partitions to the condensate interface rather than the interior in all systems examined (**Figures 3D; S12**), and filament nucleation initiates at this liquid-liquid interface (**Figures 3C; S4; S9; and Video S2**). This interfacial partitioning likely reflects a balance of electrostatic interactions between tubulin, peptide, and NA components, creating a favorable site for MT nucleation. We speculate that this arises from the unique conformational ensemble of the protein molecules at the condensate interface relative to the dilute or dense phase^81^, which may allow the interface to serve as a critical site for biochemical reactions^26, 82^. Thus, the interface appears to serve as a spatially distinct nucleation site, where local concentration enhancement, ion partitioning, and unique orientation of tubulin subunits may work together to facilitate MT nucleation, consistent with other condensate systems that show similar behavior^83^. However, our experiments cannot rule out whether tubulin and MT filaments are primarily localized at the outside or inside surface of the condensate, and future experiments with higher spatial resolution will be required.

Interestingly, the extent of tubulin enrichment, whether at the interface or within the interior, does not show a clear correlation with condensate viscoelasticity or MT assembly efficiency. This suggests that while interfaces seed MT nucleation, subsequent filament growth is constrained by the rheological properties of the condensate, which govern lateral tubulin diffusion, viscoelasticity, and bending rigidity of the interface^84^. Indeed, tubulin mobility both within and at the interface of the host condensate emerges as a key determinant of the observed MT assembly outcomes (**Figures 3F & 4E**). This is in agreement with recent studies on condensate-aided actin filament assembly utilizing VASP condensates, which revealed that condensate fluidity was important for condensate deformation and ensuing bundling of actin filaments^37, 40^. An additional regulatory force in condensate-mediated MT assembly could emerge from condensates preferentially binding to MT filaments, rather than tubulin enrichment per se, analogous to mechanisms described for actin assembly in other condensate systems^40^. Overall, our findings support a view of condensates as heterogeneous microenvironments in which physical and mechanical features cooperate to shape reaction dynamics and trajectories.

Through computer simulations, we propose a chemo-mechanical model in which a single condensate interfacial viscosity parameter simultaneously controls MT assembly through two coupled mechanisms: (i) Lateral surface diffusion of tubulin governs whether growing surface-bound tips are resupplied fast enough to sustain elongation, and (ii) Interfacial mobility is also inversely related to activation energy of dissociation for MTs which together with length-dependent elastic stress sets the lifetime of surface contact before radial escape (**Figure 5C**). Condensates with softer interface (KGKGG-type) with weakly adhesive interactions emerge as efficient catalysts for MT assembly because they can support rapid delivery of tubulin to growing MT filament tips and release filaments into the dilute phase to form asters, while highly viscoelastic condensates (RGRGG-type) with more cohesive interfaces trap and stall MT nuclei (**Figure 5F**). This framework reconciles the apparent paradox of condensates acting both as promoters and inhibitors of polymerization reactions, depending on their emergent material state.

In a broader sense, our findings hint at a defined molecular grammar linking condensate mechanics to cytoskeletal function. The ability to tune MT assembly by modulating condensate viscoelasticity and interfacial organization underscores that condensates are dynamic regulatory hubs. By connecting synthetic design principles to cytoskeletal organization, this work suggests that cellular condensates, such as centrosomes or TPX2 bodies, may exploit a similar coupling between mechanical properties and reaction dynamics to modulate MT network formation during mitosis, transport, and morphogenesis. This chemo-mechanical framework further provides design rules for engineering cytoskeletal regulation into synthetic condensates, guiding the development of programmable cytoskeletal scaffolds for artificial cells.

## Material and Methods

### Peptide and DNA samples

The peptides used in this study ([RGRGG]_5_, [KGKGG]_5_, [KGYGG]_5_, [RGPGG]_5_, and [RPRPP]_5_) were synthesized by GenScript USA Inc. (NJ, USA, >90% purity). All peptides contained a cysteine residue at the C-terminus for site-specific fluorescence labeling. Peptides were reconstituted in nuclease-free water (Santa Cruz Biotechnology) containing 50 mM dithiothreitol (DTT) to prevent cysteine oxidation. The list of peptides used in this study is as follows:

(KGKGG)_5_ = KGKGGKGKGGKGKGGKGKGGKGKGGC

(RGRGG)_5_ = RGRGGRGRGGRGRGGRGRGGRGRGGC

(RPRPP)_5_ = RPRPPRPRPPRPRPPRPRPPRPRPPC

(KGYGG)_5_ = KGYGGKGYGGKGYGGKGYGGKGYGGC

ssDNA oligos of various lengths [dT20, dT40, dT90, dT200, d(TTAGGG)_10_], and ssRNA oligo r(UUAGGG)_10,_ were purchased from Integrated DNA Technologies (IDT) and reconstituted in nuclease-free water. The reconstituted peptide and NA stock solutions were centrifuged at 23000×g for 2 minutes, and the supernatant was extracted. We confirmed the absence of microscale aggregates in all stock solutions by light microscopy, after which they were split into multiple aliquots and stored at −20 °C for further use.

### Peptide-NA condensate preparation

Condensates were prepared by mixing appropriate amounts of the peptide and ssDNA/RNA to a final concentration of 2.5 mg/mL peptide and 1.25 mg/mL ssDNA/RNA in a buffer containing 60 mM PIPES (pH 6.9), 1.5 mM MgCl_2_, 0.38 mM EGTA, and 12 mM DTT. The mass ratio of NA to the peptide is 0.5:1 unless otherwise noted. This ratio was chosen based on turbidity measurements indicating that the 0.5:1 ratio yields near-optimal phase separation across all systems tested^21, 22^. Due to the comparative nature of this study, all buffer conditions were kept identical.

### Bioinformatics analysis

Approximately 1092 human MAPs were identified using the Gene Ontology (GO) term “microtubule binding” (GO:0008017) with the taxon filter *Homo sapiens* via the QuickGO web interface^85^. Corresponding UniProt identifiers were retrieved, and full protein sequences were obtained using the UniProt API through a custom Python workflow. Intrinsically disordered regions (IDRs) were identified from UniProt feature annotations labeled “region of interest” with the description “Disordered.” Each of these “Disordered” segments was validated by an independent disorder-prediction tool, IUPred3^86^. The start and end coordinates of these annotated regions were used to extract the corresponding sequence segments from each protein sequence. To obtain reference amino acid frequencies for the human proteome, UniProt entries for proteins (organism_id: 9606) were retrieved using the UniProt API and parsed using a separate custom Python script to tabulate amino acid frequencies. A third Python script was used to analyze amino acid composition in the resulting datasets. Amino-acid occurrences were counted across all extracted MAP IDR sequences, excluding ambiguous residues (denoted as “X” in UniProt entries). Counts were normalized by the total number of residues to obtain relative amino acid frequencies for MAP IDRs. These frequencies were compared with reference amino acid frequencies for the human proteome. Enrichment for each residue was calculated as the log_2_ ratio of the observed frequency in MAP IDRs to the corresponding frequency in the human proteome. The resulting enrichment values were visualized as a heatmap (**Figure 1A**).

### Turbidity measurements

Samples were prepared in a tube by mixing the peptide and the ssDNA at the desired mixing ratio and a fixed peptide concentration of 2.5 mg/mL. The buffer of these samples contains 60 mM PIPES (pH 6.9), 1.5 mM MgCl_2_, 0.38 mM EGTA, and 12 mM DTT. The sample was then placed on a UV-Vis spectrophotometer (NanoDrop 1C) and the solution turbidity at 350 nm was measured for three independent samples. Before measuring the turbidity of peptide-ssDNA samples, the instrument was blanked using the experimental buffer. The turbidity values were averaged for each mixing ratio, and the error was estimated as the standard deviation (SD) from three replicates.

### Coverslip, and chamber slide well preparation

For all experiments in this work, Tween20 coating was used to prevent droplet spreading on the glass surface. Coverslips and chamber slide wells were first cleaned by incubation in 2% (v/v) Hellmanex solution for 2 hr, rinsed 5-6 times with MilliQ water. Next, they were immersed in a 20% (v/v) Tween20 solution for 30 min. Subsequently, the slides and coverslips were rinsed 6–7 times with MilliQ water and dried using compressed air. Finally, the slides and coverslips were dried in an oven set at 37 °C for 16 hr and stored at room temperature for later use.

### Tubulin partitioning and MT assembly assay

Tubulin and HiLyte 647-labeled tubulin were purchased from Cytoskeleton Inc. The synthetic condensate MT polymerization assay was carried out based on methods described previously^50, 87^. Briefly, condensates were formed in a buffer containing 60 mM PIPES (pH 6.9), 1.5 mM MgCl_2_, 0.38 mM EGTA, and 12 mM DTT and allowed to equilibrate within a microfuge tube or in a chamber slide well for 20 min, before the addition of a 10 μM tubulin and 2 mM GTP without mixing (see also, **Figure S3**). Unlabeled tubulin was doped with 900 nM ^Hy647^Tubulin for visualization by fluorescence microscopy. For the tubulin partitioning experiments, only ^Hy647^Tubulin is introduced to the condensate sample mixture at a concentration specified in the text. The final sample was sandwiched between a Tween-20-coated coverslip and a glass slide, separated by three layers of double-sided tape. The remaining void in the sandwich was filled with mineral oil to prevent evaporation. For time-lapse fluorescence imaging, a sample was applied to chamber slide wells with lids (Grace Biolabs). Q2 laser scanning confocal microscope (ISS Inc., 63x objective) was used for fluorescence imaging.

Fluorescence images collected for partition coefficient image analysis were analyzed using Fiji^88^ to segment individual condensates and the mean intensity of tubulin was estimated at the interface and the interior, and the partition coefficient was calculated as either the ratio of intensity value at the interface of condensates (interface) or within condensates (interior), divided by the average dilute phase intensity determined from at least 10 randomly chosen regions without condensates. Plots of partition coefficients show the median and standard deviation of measurements from at least 150 condensates across three independently prepared replicates.

### Image analysis and quantification of MT coverage, and MT filament length

Confocal fluorescence images visualized with ^Hy647^Tubulin were used as input to estimate MT surface coverage and MT filament length/width. Confocal fluorescence images visualized with AF488-peptide were used as input for estimating condensate surface coverage. Random frames were imaged to ensure an accurate representation of the surface area.

Binary masks for MTs were generated by ridge-based filament detection followed by dilation to the diffraction-limited filament width. Because imaging resolution limited precise estimation of filament width, MT coverage was quantified as an effective area occupied by diffraction-limited filaments using a consistent ridge-detection–based approach across all conditions. Condensates were segmented independently by Huang2 automated intensity thresholding. Using the ‘Analyze Particles’ function of Fiji, masks of individual condensate (sizes > 0.5μm^2^; ellipticity 0.5-1.0) were determined. The MT area outside condensates was computed by logical intersection of MT and inverted condensate masks. Subsequently, the area fractions occupied by MT filaments or condensates were normalized to the total area of the image acquisition (corresponding to the tubulin and peptide channels) and estimated.

Ridge detection was performed using the ImageJ Ridge Detection plugin^89^, which implements the unbiased curvilinear structure detector^90^, to quantify MT filament length and width (**Figure S7**) across different mixing ratios, by keeping constant input parameters across all samples. For each mixing ratio, around 3000-5000 filaments were quantified from at least 3 replicates. The plugin seemed to underestimate the number of detected filaments.

### FRAP measurements

FRAP measurements were carried out on the FRAP module installed on LAS X software, using a STELLARIS 5 confocal microscope configured on a DMi8 inverted microscope (Leica). Samples were prepared in 60 mM PIPES (pH 6.9), 1.5 mM MgCl_2_, 0.38 mM EGTA, and 12 mM DTT at the indicated peptide and NA concentrations with the addition of 900 nM ^Hy647^Tubulin. Bleaching was performed on a circular ROI ∼1 μm in diameter with 100 % laser power at peak excitation wavelength for the given fluorophore to result in bleaching to ∼20 % of prebleached fluorescence intensity. Recovery of the bleached region was monitored every 0.378 s for a total of 240 s. Intensity values were normalized by dividing by values of an unbleached condensate in the same field of view to account for general photobleaching and then dividing by the resulting maximum value to set the highest intensity to 1. Plots were generated in GraphPad Prism 10 and represent means and SDs from 3-7 FRAP curves across two independent replicates (**Figures 3E; 4F**). The FRAP recovery curves (**Figure S11**) were also plotted using IgorPro6.23, and the best fit was obtained using 𝐼(𝑡) = 𝑎 + b (𝑡/𝜏 ½)/ 1 + (𝑡/𝜏 ½) as reported previously^91, 92^. All datasets produced a good fit for this model. Here 𝐼(𝑡) is the normalized and corrected relative intensities over time; 𝑎, and 𝑏 are the normalized intensity in the bleach ROI immediately following bleach, and after full recovery (plateau), respectively; and 𝜏 ½ denotes recovery half-times. The mobile fractions (𝑀𝑓) were also calculated from the data obtained from normalized recovery curves using the equation 𝑀𝑓 = [(𝑏 − 𝑎)/(𝑐 − 𝑎)] ∗ 100 as previously described^92^, where 𝑎, 𝑏, and 𝑐 are the normalized intensities in the bleach ROI immediately following bleach, after full recovery (plateau), and before the bleach, respectively.

### DLS measurements to estimate cluster size

The size distribution was measured using dynamic light scattering (Lite Sizer 500, Anton Paar). Samples were prepared in a tube by mixing the peptide and the ssDNA at the desired mixing ratio and a fixed peptide concentration of 2.5 mg/mL. The buffer for these samples contains 60 mM PIPES (pH 6.9), 1.5 mM MgCl_2_, 0.38 mM EGTA, and 12 mM DTT, and the total sample volume was 50 µL. The measurements were initiated after 15 min of sample preparation by loading the sample into a low-volume cuvette (Univette) and measuring at 25 °C using side scatter (automatic angle). The sizes of the condensates were obtained using the General analysis mode and the Advanced cumulant model^93^. The measurements were repeated twice, and the average values are reported (**Figure 2D**).

### Viscosity estimation from Video Particle Tracking (VPT) nanorheology

To estimate the viscosity of the peptide-ssDNA condensates, we prepared the samples as described in the “Peptide-NA condensate preparation section”, except we added yellow-green, fluorescent carboxylate-modified 200-nm beads (Invitrogen) as probe particles before inducing phase separation. The prepared samples were placed on coverslips coated with 20% Tween20 (v/v) solution and were sandwiched with a glass slide using double-sided tape. The samples were then taken to the Zeiss Axio Observer inverted microscope equipped with a 100× oil immersion objective. The samples were allowed to rest on the microscope until no fusion events were observed, and the droplets were settled on the coverslip surface. The videos of the bead motion inside the condensates were acquired using a back-illuminated Kinetix 22 sCMOS camera (Teledyne Photometrics) with an exposure time of 100 ms for 1000 frames. These videos were used to extract the trajectories in two dimensions for each tracked bead inside the condensate using the TrackMate plugin of Fiji software^94, 95^. The quality and intensity filters in TrackMate were used to avoid tracking aggregated beads.

The details of the analysis performed to obtain the MSD of the beads are described in our previous work^7^. Briefly, the extracted trajectories were corrected for drifting by subtracting the center of mass trajectory, which was calculated using the velocities of particles in the following way

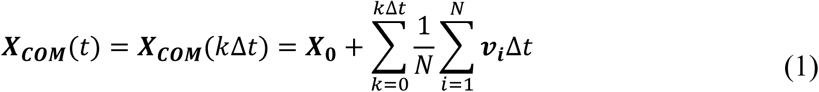

Where 𝑘 is the frame number, Δ𝑡 is the frame time, and 𝑁 is the number of particles per frame. X_𝟎_is the center of the mass vector of the first frame. After correcting the trajectories for drifting, the mean squared displacement (MSD) was calculated as

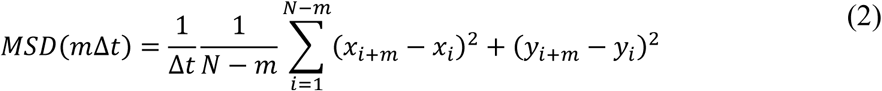

Where 𝑚 is the lag time in frames. The MSD was then fitted using^57, 96^

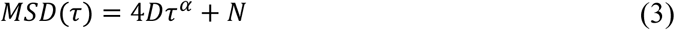

To obtain the diffusion coefficient 𝐷 of the particles. For all the systems, we ensured that the value of the diffusivity exponent 𝛼 is equal to 1 by choosing an appropriately long frame time to ensure measuring the terminal viscous behavior. The diffusion coefficient is then converted to viscosity using the Stokes-Einstein equation^57, 96^

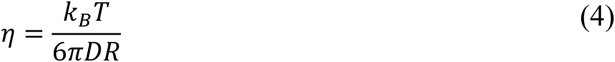

where 𝑅 is the radius of the particles and 𝑇 is the temperature of the sample. The mean viscosity and the standard deviation from the mean are reported from VPT measurements on 12-15 condensates over 3 independently prepared sample replicates. We note that, due to differences in buffer and pH conditions, the viscosity of arginine-based condensates is higher than previously reported ^7, 21, 22^.

### Dextran mesh size determination

Peptide-NA samples were prepared at 5.0 mg/mL peptide and 2.5 mg/mL dT40 DNA in a buffer containing 60 mM PIPES (pH 6.9), 1.5 mM MgCl_2_, 0.38 mM EGTA, and 12 mM DTT. After mixing the peptide and the NA and forming condensates, a small concentration (∼500 nM) of tetramethylrhodamine isothiocyanate (TRITC)-labeled dextran of the desired size was added to the sample. The sample was mixed and subsequently imaged under a confocal microscope (LUMICKS C-trap). If the hydrodynamic radius of the dextran molecule is smaller than the mesh size of the condensate, it is expected that dextran molecules will be positively recruited into the condensates. However, if the hydrodynamic radius of the dextran molecule is larger than the mesh size of the condensates, dextran recruitment will not occur. The partition behavior of dextrans of variable molecular weights (and variable hydrodynamic radii) was recorded and the mesh size range was determined accordingly^21, 67^. The smallest dextran molecule that partitions negatively into the condensate defines the upper limit of the mesh size.

### Software

Fiji^88^ (version 1.54f) was used for processing and analyzing microscopy images. Custom python scripts were used for bioinformatic analysis of MAP IDRs. A custom Python script was used for plotting VPT nanorheology data and estimation of dynamical moduli. The scripts are available publicly (*see* ‘Code Availability’ section). Trackmate (v7.14) was used for nanorheology-related analysis. GraphPad Prism 10 (v10.4.1) was used for all the plots except FRAP recovery traces, which used IgorPro6.23 for fitting analysis. Biorender (2025) was used for preparing schematics (**Figures 3C; D; and 5F**), while Adobe Illustrator CC(2023, v23.0) was used for figure assembly and production. LAS X software was used for acquiring fluorescence images with the DMi8 inverted microscope (Leica). Vistavision (v4.2) was used for acquiring fluorescence images with the ISS Q2 laser scanning confocal microscope.

### Computer simulations

We developed and employed a coarse-grained stochastic model of MT nucleation, growth, and detachment at the surface of a spherical condensate droplet. The details of simulations, including equations of reaction-diffusion steps and parameters used, are provided in the Supplementary Information. All the scripts used to produce the data reported in the manuscript are deposited in GitHub, https://github.com/PotoyanGroup/MT-interface-growth

## Acknowledgements

This work was supported by the US National Institutes of Health (NIH) through grants R35 GM138186 (to P.R.B.), R35 GM138243 (to D.A.P), the St. Jude Children’s Research Collaborative on the Biology and Biophysics of RNP Granules (to P.R.B.), and F32GM161082 (to S.S). We would like to acknowledge Shamli Manasvi for help with the bioinformatics analysis, Tirth Bhatta for the brightfield and turbidity measurements, and Joe Basalla for technical support with FRAP measurements. We would like to thank Dr. Sangwoo Shin for sharing the Lite Sizer 500 instrument. Also, the authors greatly appreciate the critical feedback from the Banerjee laboratory’s group members. Any opinions, findings, conclusions, or recommendations expressed in this material are those of the author(s) and do not necessarily reflect the views of the NSF, NIH, or any other bodies.

## Author Contributions

Conceptualization: P.R.B. and S.S.; Methodology: S.S, A.S., D.A.P; Investigation: S.S., A.S., D.A.P; Data curation: P.R.B., S.S., A.S., D.A.P; Formal analysis: S.S., A.S., D.A.P; Visualization: P.R.B., S.S., A.S., D.A.P; Writing – original draft: P.R.B. and S.S.; Writing – review & editing: all authors; Supervision: P.R.B.; Funding acquisition: P.R.B., D.A.P, and S.S.

## Data Availability

Unless otherwise stated, all data supporting the results of this study can be found in the article and the supplementary files.

## Code Availability

Codes for nanorheology are available via GitHub at https://github.com/BanerjeeLab-repertoire/Synthetic-Biomolecular-Condensates-as-Tunable-Microtubule-Assembly-Hubs.

## Declaration of Generative AI and AI-assisted technologies

The authors acknowledge the use of ChatGPT5.3 and Microsoft Copilot to enhance the readability and clarity of certain sections of the manuscript text. The authors reviewed and edited all content generated with the assistance of AI tools and take full responsibility for the content of this manuscript.

