## Supplementary File for "Synthetic Biomolecular Condensates as Tunable Microtubule Assembly Hubs"

### Supplementary Figures

A

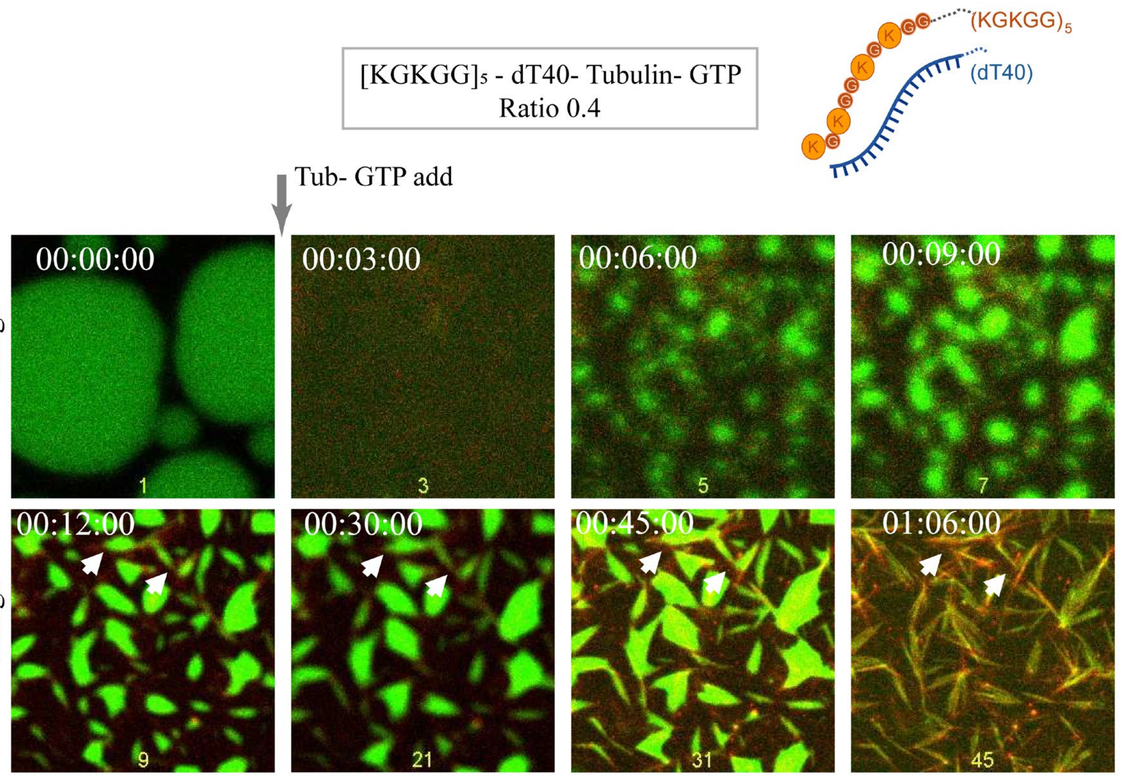


B

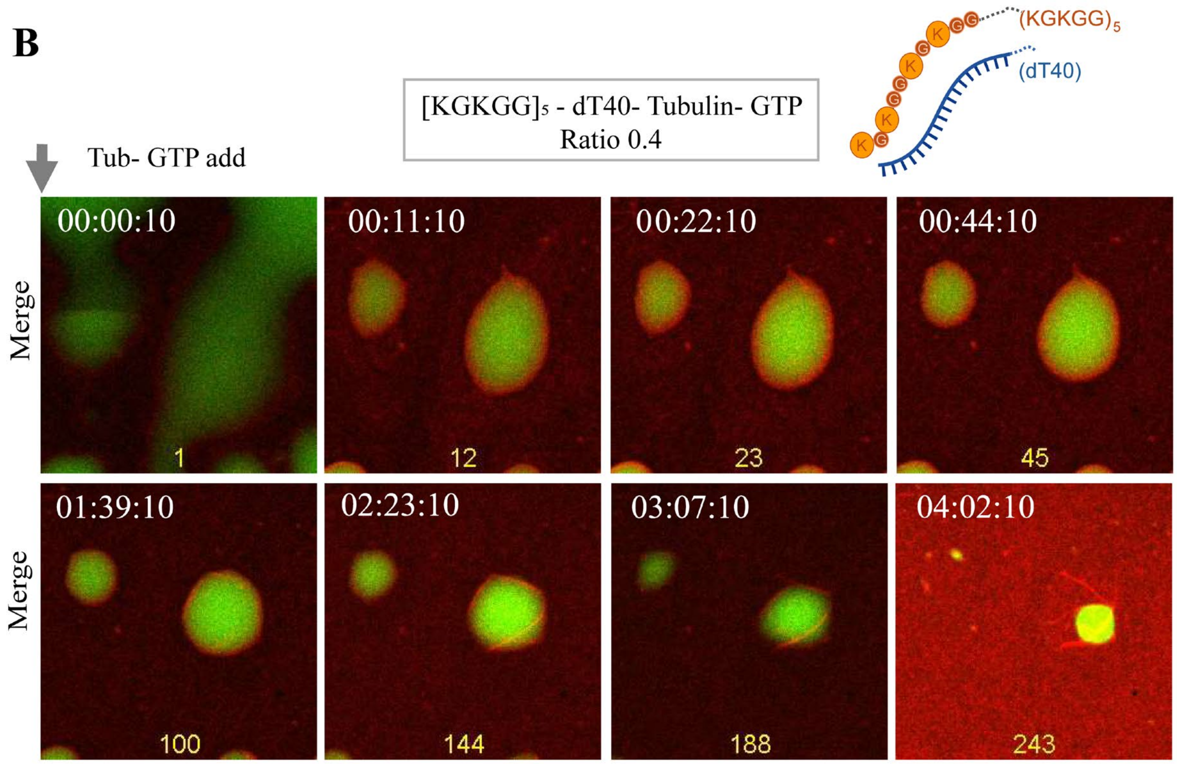

**Figure S1. Synthetic peptide-NA condensates catalyze microtubule (MT) assembly. (A)** Time-lapse images correspond to Video S1. Addition of 10 μM tubulin doped with 900 nM $^{Hy647}$Tubulin and 2 mM GTP to $[KGKGG]_5$–dT40 condensates containing 500 nM $^{AF488}[KGKGG]_5$. The microscope objective was focused on a suitable position in the middle of the sample with droplets. The VistaVision software (ISS Inc.) was set to acquire time-lapse images continuously every 90 s before tubulin-GTP (Tub-GTP) addition. Once imaging is started, a 1.0-1.2 μl volume of Tub-GTP mix was pipetted on the sample close to the image acquisition spot. Panels are visualized as an overlay of $^{Hy647}$Tubulin & $^{AF488}[KGKGG]_5$ fluorescence, with panel labelled 1 (timestamp 00:00:00) showing $[KGKGG]_5$–dT40 condensates before Tub-GTP addition, and subsequent panels labelled 3-45 (timestamps 00:03:00 to 01:06:00) visualized immediately after Tub-GTP addition. The white arrows highlight representative regions showing droplet fusion and wetting on MT filament projections and eventual filament branching. **(B)** Time-lapse images correspond to Video S2. Addition of 5 μM tubulin doped with 500 nM $^{Hy647}$Tubulin and 1 mM GTP to $[KGKGG]_5$–dT40 condensates doped with 500 nM $^{AF488}[KGKGG]_5$. The objective was focused on a suitable position in the middle of the sample with droplets. A 1.0-1.2 μL volume of Tub-GTP mix was added to the sample using a pipette far from the image acquisition spot. The VistaVision software (ISS Inc.) was set to acquire time-lapse images continuously every 60 s immediately following Tub-GTP addition. Panels are visualized as an overlay of $^{Hy647}$Tubulin & $^{AF488}[KGKGG]_5$ fluorescence immediately after Tub-GTP addition showing MT nucleation and growth at the interface of a condensate and subsequent shrinkage of the condensate with filament growth (timestamps 00:00:10 to 03:59:10). For A and B, the condensate composition and sample conditions are as reported in Videos S1 and S2. Scale bar is 5 μm.

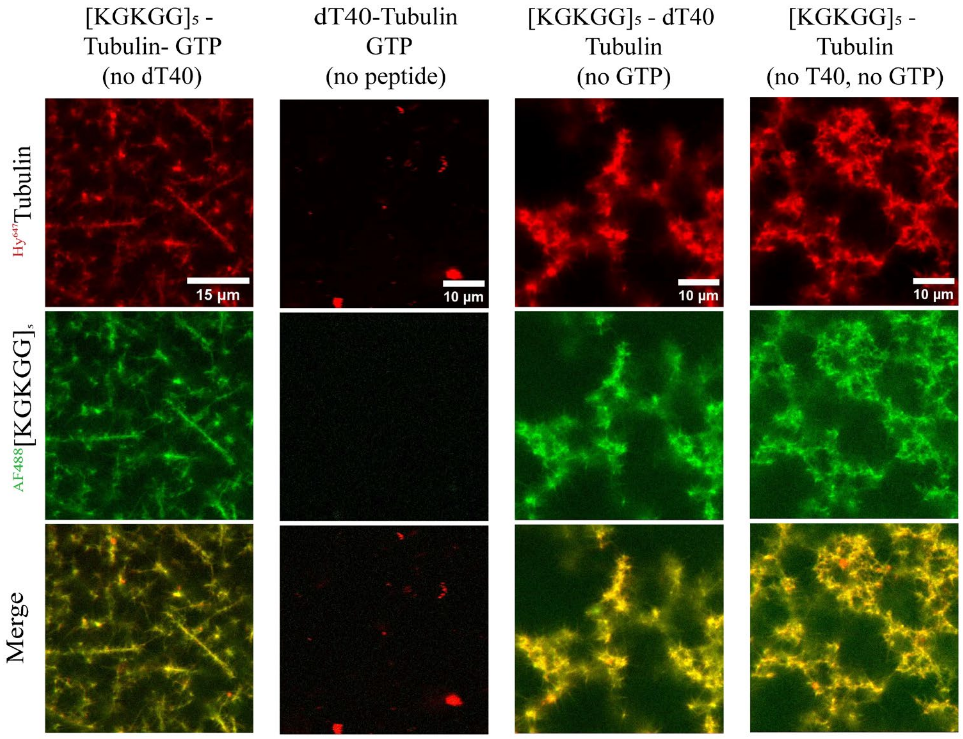


**Figure S2. Robust MT assembly requires all four components in this multi-component system.** Representative confocal fluorescence micrographs of three-component systems lacking the fourth component: dT40, peptide or GTP (first 3 panels from the left); and a two-component system lacking: dT40, and GTP ($4^{th}$ panel on right), with 10 μM tubulin (red, top), with or without 2 mM GTP, with or without 2.5 mg/mL $[KGKGG]_5$ (green, middle), 1.25 mg/mL dT40 and overlay (merge, bottom). 500 nM Alexa-fluorophore-labeled peptides and 900 nM $^{Hy647}$Tubulin were used for imaging, as indicated. These samples do not support robust MT filament assembly. Samples were imaged 60 min after tubulin addition. Scale bar is 5 μm.

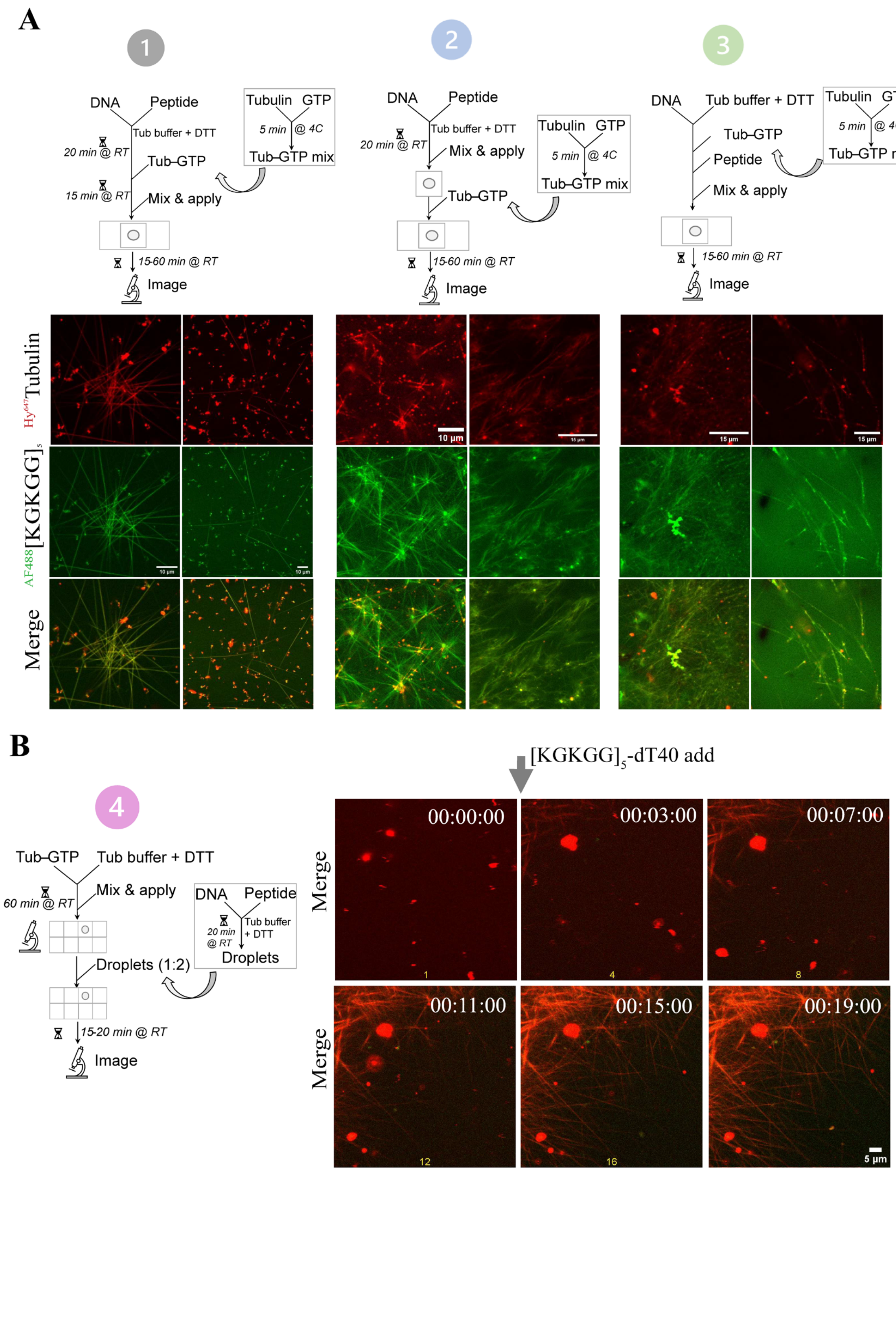
A
1
DNA
Peptide
Tub buffer + DTT
20 min @ RT
Tub–GTP
15 min @ RT
Mix & apply
Tubulin GTP
5 min @ 4C
Tub–GTP mix
15-60 min @ RT
Image
2
DNA
Peptide
Tub buffer + DTT
20 min @ RT
Mix & apply
Tub–GTP
Tubulin GTP
5 min @ 4C
Tub–GTP mix
15-60 min @ RT
Image
3
DNA
Tub buffer + DTT
Tub–GTP
Peptide
Mix & apply
Tubulin GTP
5 min @ 4C
Tub–GTP mix
15-60 min @ RT
Image
Hy647Tubulin
AF488[KGKGG]5
Merge
10 μm
15 μm
B
4
Tub–GTP
Tub buffer + DTT
60 min @ RT
Mix & apply
DNA
Peptide
20 min @ RT
Tub buffer + DTT
Droplets
Droplets (1:2)
15-20 min @ RT
Image
[KGKGG]5-dT40 add
Merge
00:00:00
00:03:00
00:07:00
00:11:00
00:15:00
00:19:00
5 μm

**Figure S3. Experimental setups involving different order of component addition show mildly variable outcomes. (A)** Panels from left to right (1, 2, 3) each represent three different experimental setups employed to track MT assembly in the presence of peptide-NA condensates. The schematic shows differences including component addition in microfuge tubes versus coverslip, order of component addition, incubation times & pipette aided mixing. In each of these scenarios, representative confocal fluorescence micrographs show organization of MT filaments, with 10 μM tubulin (red, top), 2 mM GTP, 2.5 mg/mL $[KGKGG]_5$ (green, middle), 1.25 mg/mL dT40 and overlay (merge, bottom). Samples imaged 60 min after preparation. The final sample contained 60 mM PIPES (pH 6.9), 1.5 mM $MgCl_2$, 0.38 mM EGTA and 12 mM DTT. All experiments in the main figures and associated supplementary figures and videos involved the setup described in scenario 2 which reproducibly showed the best MT assembly outcomes. **(B)** The schematic on the left, shows a 4$^{th}$ scenario (experimental setup) where a tubulin-GTP mixture was formed in the presence of 25 μM tubulin, 2 mM GTP, 900 nM $^{Hy647}$Tubulin, 60 mM PIPES (pH 6.9), 1.5 mM $MgCl_2$, 0.38 mM EGTA and 12 mM DTT, following which pre-formed $[KGKGG]_5$–dT40 condensates (2.5 mg/mL $[KGKGG]_5$, 1.25 mg/mL dT40, 500 nM $^{AF488}[KGKGG]_5$ in the same buffer) were added (1:2) without additional mixing. Notably, prior to condensate addition, despite the higher tubulin concentrations, tubulin did not spontaneously self-assemble into long filaments under these conditions. On the right, time-lapse images captured before and after condensate addition. Panels are visualized as an overlay of $^{Hy647}$Tubulin & $^{AF488}[KGKGG]_5$ fluorescence showing MT nucleation and growth immediately after condensate addition (timestamps 00:03:00 to 00:19:00). Scale bar is 5 μm.

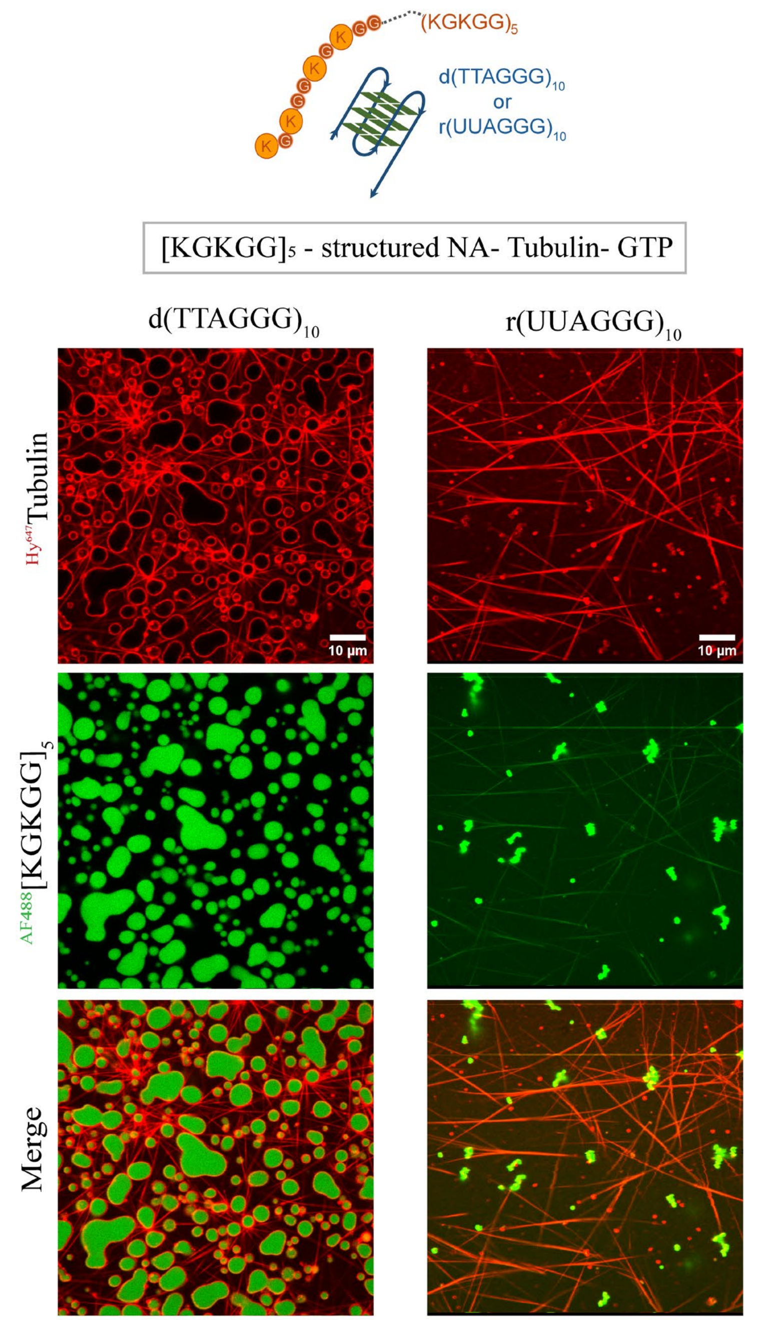


**Figure S4. Effect of NA sequence and structure on condensate-mediated MT assembly.** Representative confocal fluorescence micrographs showing organization of MT filaments in the presence of $[KGKGG]_5$–$d(TTAGGG)_{10}$ condensates (left panels), and $[KGKGG]_5$–$r(UUAGGG)_{10}$ condensates (right panels). The NA components comprised 10 repeats of (TTAGGG), and (UUAGGG), respectively, which are known to adopt stable G-quadruplex structures[1-4]. Samples with 10 μM tubulin (red, top), 2 mM GTP, 2.5 mg/mL $[KGKGG]_5$ (green, middle), 1.5 mg/mL DNA or 1.0 mg/mL RNA and overlay (merge, bottom). 500 nM Alexa-fluorophore-labeled peptides and 900 nM $^{Hy647}$Tubulin are used for imaging, as indicated. Samples were imaged 60 min after tubulin-GTP addition.

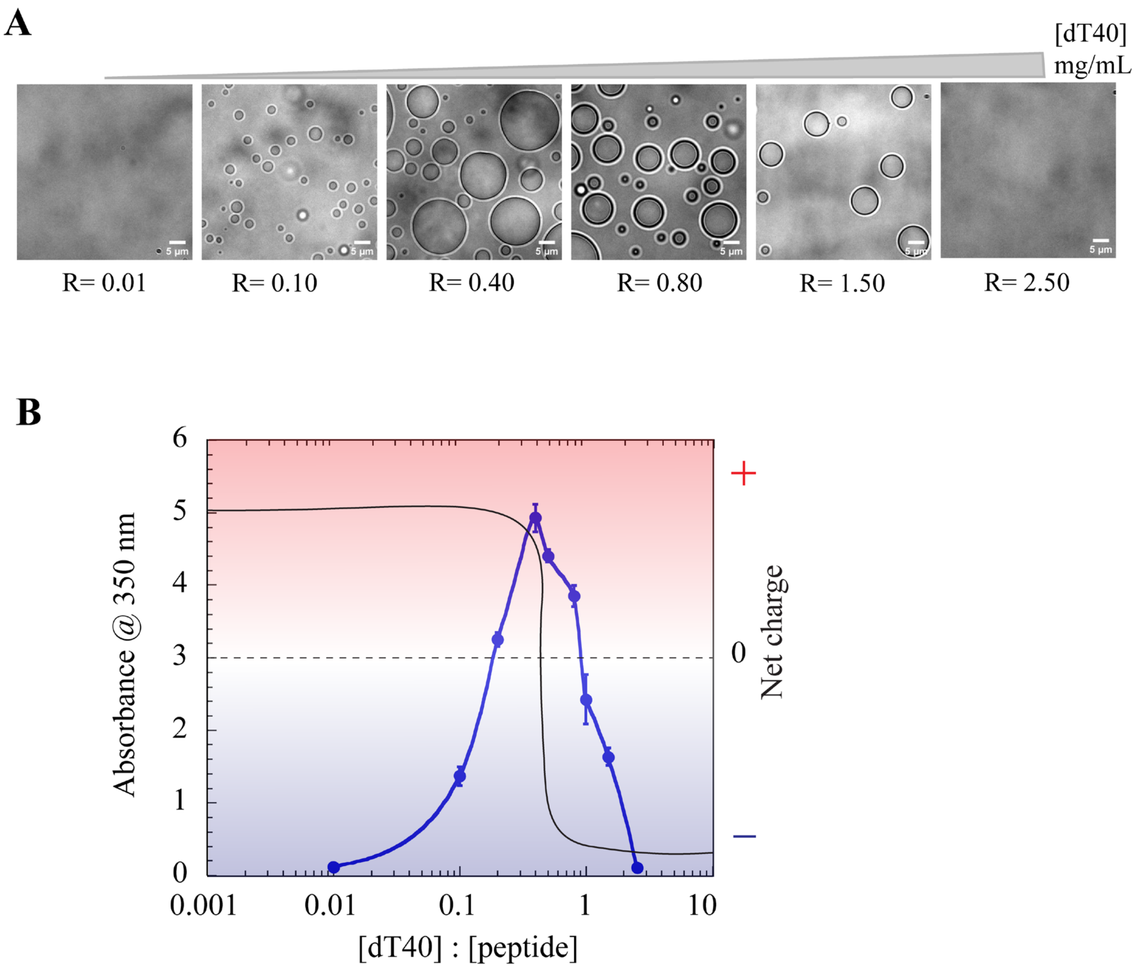


**Figure S5. Condensate formation by [KGKGG]$_5$-dT40 mixtures at different [dT40]:[peptide] ratios show reentrant phase behavior and charge inversion. (A)** Bright-field images of [KGKGG]$_5$-dT40 condensates with increasing [dT40] concentrations. The [KGKGG]$_5$ was fixed at 2.5 mg/mL, while [dT40] varied between 0.03 - 6.3 mg/mL. **(B)** Solution turbidity at 350 nm (left axis, blue points and line) as a function of increasing [dT40]-to-[peptide] ratios. The blue line is drawn as a guide to the eye. The incubation time for this measurement is 2 - 3 minutes after vigorous mixing. The data shows three independent measurements with standard deviation reported. The composition of the condensate systems is 2.5 mg/mL [KGKGG]$_5$ and 0.03 - 6.25 mg/mL dT40, with the final sample containing 60 mM PIPES (pH 6.9), 1.5 mM $MgCl_2$, 0.38 mM EGTA and 12 mM DTT. The turbidity plot is overlayed with a schematic of electrophoretic mobility for [KGKGG]$_5$-NA condensates showing charge inversion in the presence of excess nucleic acid[5] (right axis, black line).

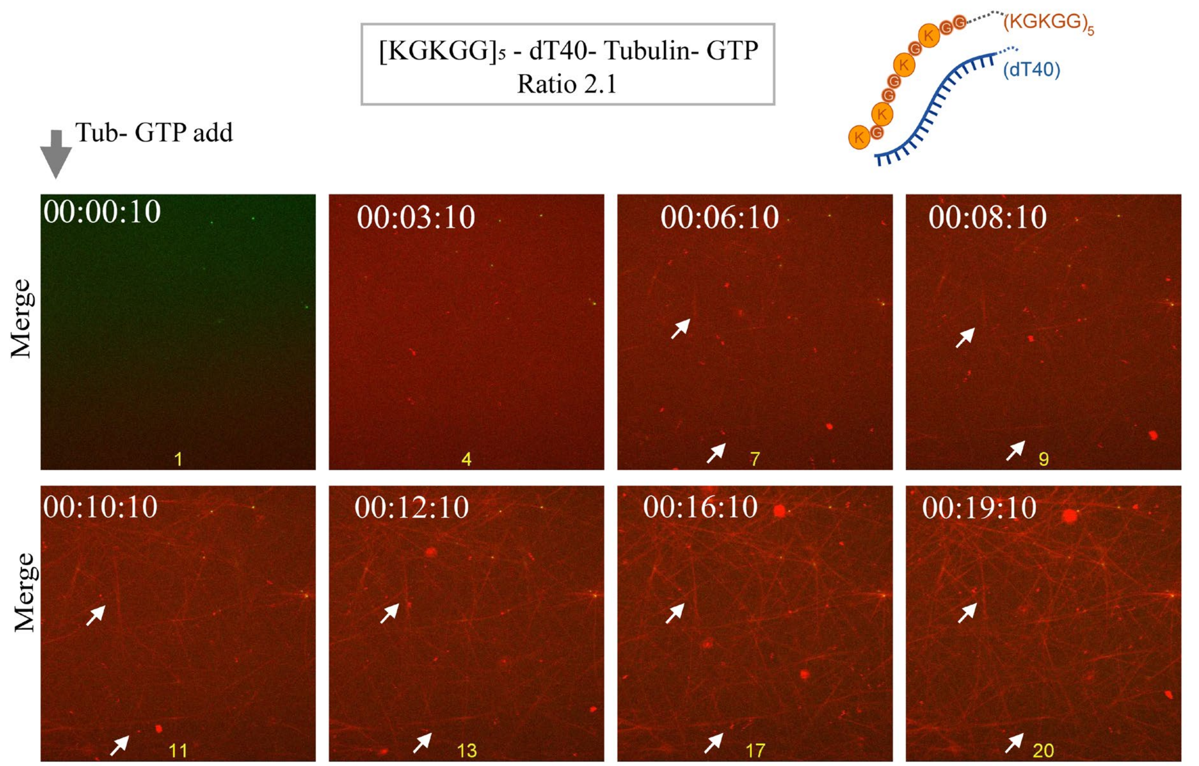


**Figure S6. Rapid MT nucleation & assembly from lysine-rich peptide-NA nanoscale complexes.** Addition of 10 µM tubulin doped with 900 nM $^{Hy647}$Tubulin and 2 mM GTP to $[KGKGG]_5$–dT40 mixture doped with 500 nM $^{AF488}[KGKGG]_5$. These observations correspond to Video S4. Panels are visualized as an overlay of $^{Hy647}$Tubulin & $^{AF488}[KGKGG]_5$ fluorescence immediately after tubulin-GTP addition, showing rapid MT nucleation and growth (timestamps 00:00:10 to 00:19:10). The white arrows highlight representative regions showing MT filaments forming in regions devoid of micron-scale condensates and eventual filament branching. Scale bar is 5 µm.

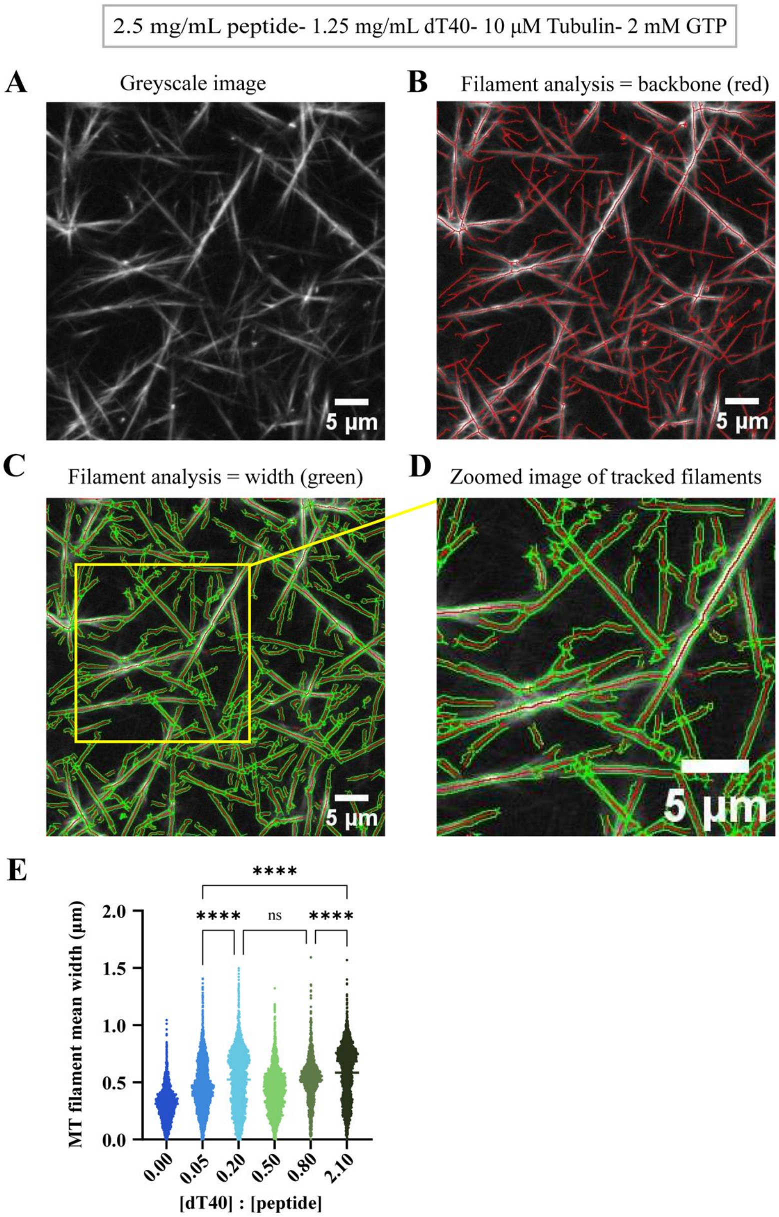


**Figure S7. Quantification of MT length and width using ImageJ. (A)** A representative greyscale image of MT filaments from Figure 2B. **(B)** MT filament analysis (length) performed on the image in panel (A) using the Ridge detection plugin[6]. Details of the image analysis protocol are described in the Methods section. **(C)** MT filament analysis (width) performed using the Ridge detection plugin. **(D)** Zoomed in region from panel C (yellow box). Image analysis allows selective segmentation of MT filaments, yielding an estimation of MT length and width. **(E)** Quantification of MT filament width corresponding to Figure 2B. Horizontal lines at each [dT40]:[peptide] ratio (R) represents the mean values. The individual data points represent measurements based on three to four independent replicates. Sample size: R = 0.00, n = 3234; R = 0.05, n = 4804; R = 0.20, n = 4777; R = 0.50, n = 4214; R = 0.80, n = 3056, and R = 2.10, n = 4665. Statistical significance was determined using an one-way ANOVA between the individual ratios ("ns" means non-significant, * means $p < 0.05$, ** means $p < 0.01$, *** means $p < 0.001$, **** means $p < 0.0001$). The associated $P$ values are shown only for R = 0.05 vs. R = 0.20 & R = 2.10; R = 0.20 vs. R = 0.80 and R = 0.80 vs. R = 2.10.

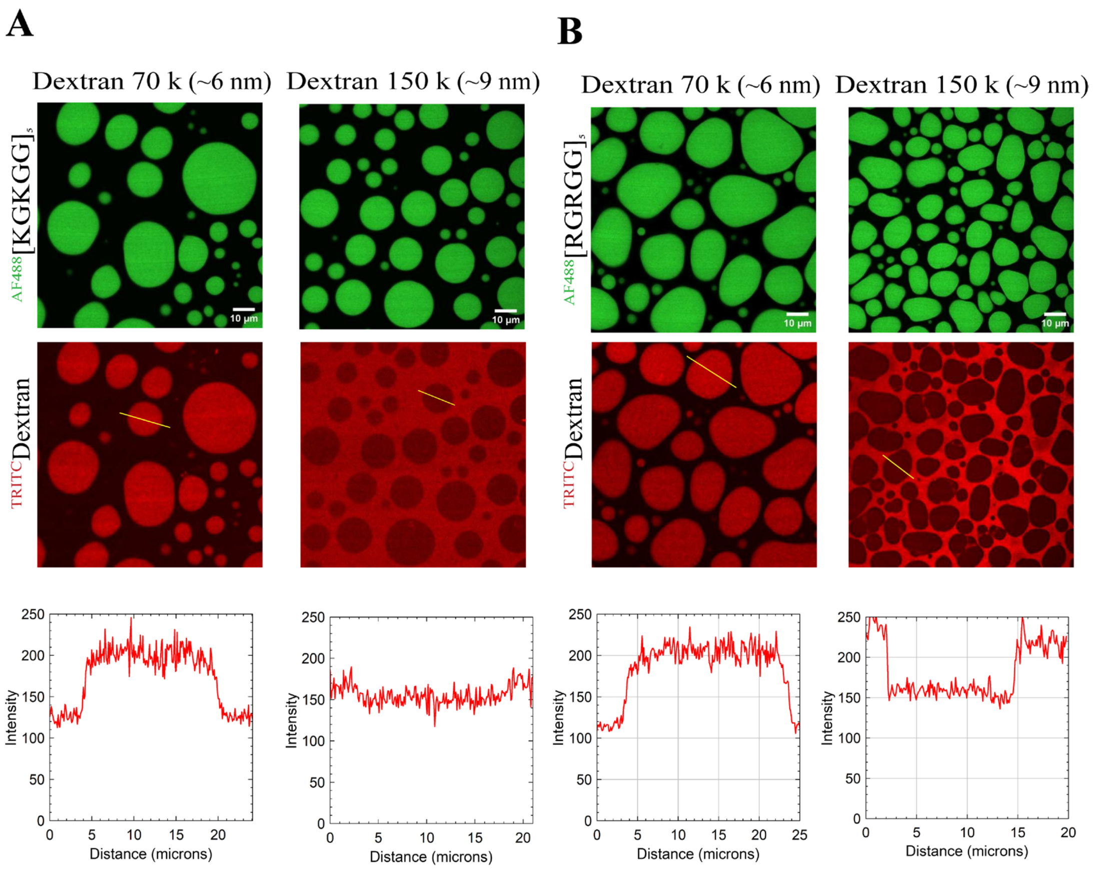


**Figure S8. Mesh size determination in [KGKGG]$_5$-dT40 and [RGRGG]$_5$-dT40 condensates using TRITC-labeled Dextran molecules with variable molecular weights** for **(A)** [KGKGG]$_5$-dT40 condensates, and **(B)** [RGRGG]$_5$-dT40 condensates. The numbers in parentheses indicate the estimated hydrodynamic radius of the Dextran molecules in aqueous solutions[7]. The upper panel shows the condensates as visualized by $^{AF488}$peptide fluorescence. The middle panel shows the partitioning of Dextran molecules within peptide-NA condensates. The scale bar represents 10 µm. The composition of the condensate systems is 2.5 mg/mL peptide and 1.25 mg/mL dT40, 500 nM $^{AF488}$peptide, 500 nM $^{TRITC}$Dextran, with the final sample containing 60 mM PIPES (pH 6.9), 1.5 mM $MgCl_2$, 0.38 mM EGTA and 12 mM DTT. The lower panel shows corresponding intensity profiles for Dextran molecules. The smallest Dextran molecule that partitioned negatively in the condensate signifies the upper limit of the mesh size. The 150k Dextran partitioned ~1.5 fold better into [KGKGG]$_5$-dT40 condensates when compared to [RGRGG]$_5$-dT40 condensates.

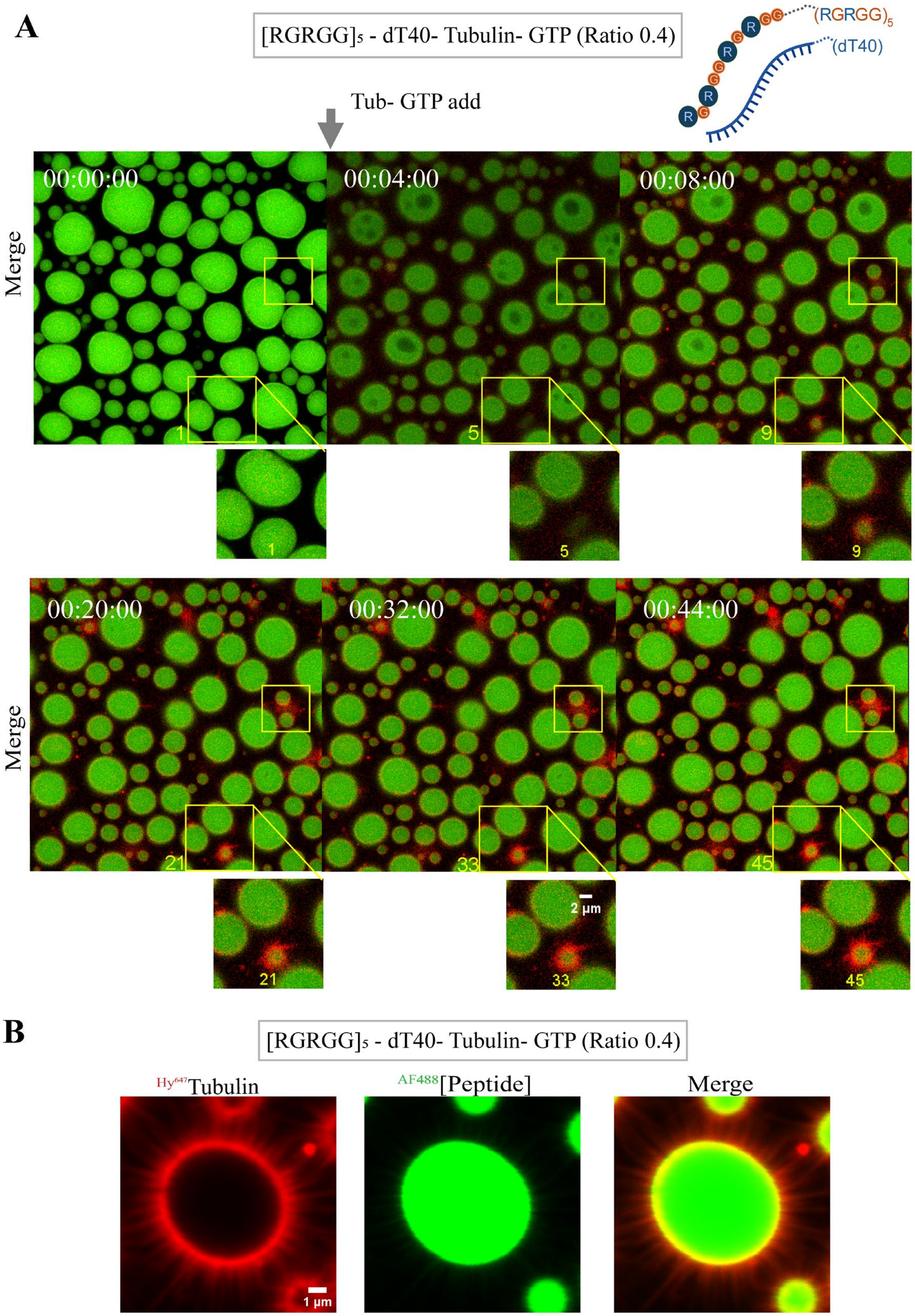
A
[RGRGG]5 - dT40- Tubulin- GTP (Ratio 0.4)
(RGRGG)5
(dT40)
Tub- GTP add
00:00:00
00:04:00
00:08:00
Merge
1
5
9
00:20:00
00:32:00
00:44:00
Merge
21
33
45
2 µm
B
[RGRGG]5 - dT40- Tubulin- GTP (Ratio 0.4)
Hy647Tubulin
AF488[Peptide]
Merge
1 µm

**Figure S9. MT assembly in the presence of arginine-rich peptide-NA condensates. (A)** Addition of 10 μM tubulin doped with 900 nM $^{Hy647}$Tubulin plus 2 mM GTP to $[RGRGG]_5$–dT40 condensates containing 500 nM $^{AF488}[RGRGG]_5$. These observations correspond to Video S5. Panels are visualized as an overlay of $^{Hy647}$Tubulin and $^{AF488}[RGRGG]_5$ fluorescence, with panel labeled 1 (timestamp 00:00:00) showing $[RGRGG]_5$–dT40 condensates before tubulin-GTP (Tub-GTP) addition, and subsequent panels labeled 5-45 (timestamps 00:05:00 to 00:45:00) visualized immediately after Tub-GTP addition. Scale bar is 5 μm. **(B)** Representative confocal fluorescence micrographs showing organization of short surface-arrested MT filaments at the interface of $[RGRGG]_5$–dT40 condensates, with 9.5 μM tubulin (red, 1st panel), 2 mM GTP, 2.5 mg/mL $[RGRGG]_5$ (green, 2nd panel), 1.00 mg/mL dT40, and overlay (merge, 3rd panel). Samples were imaged 60 min after tubulin-GTP addition.

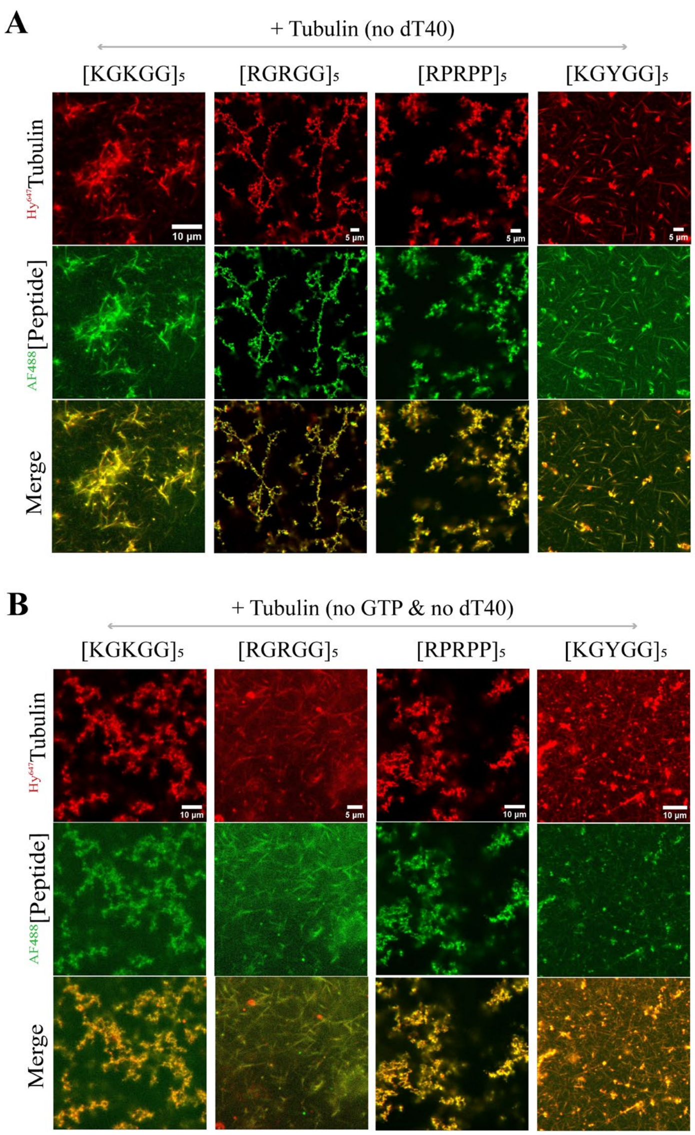


**Figure S10. Differential interaction of multivalent peptides with tubulin in the presence and absence of GTP.** Representative confocal fluorescence micrographs of three-component systems lacking the fourth component: dT40 **(A)**; and a two-component system lacking: dT40, and GTP **(B)**; with 10 µM tubulin (red, top), with or without 2 mM GTP, with 2.5 mg/mL peptide (green, middle), and overlay (merge, bottom). 500 nM Alexa-fluorophore-labeled peptides and 900 nM $^{Hy647}$Tubulin are used for imaging, as indicated. These samples do not support robust MT filament assembly, but the lysine-rich peptides form short MTs in the presence of tubulin irrespective of the presence of GTP, while the arginine-rich peptides form aggregates with tubulin. Samples were imaged 60 min after tubulin addition.

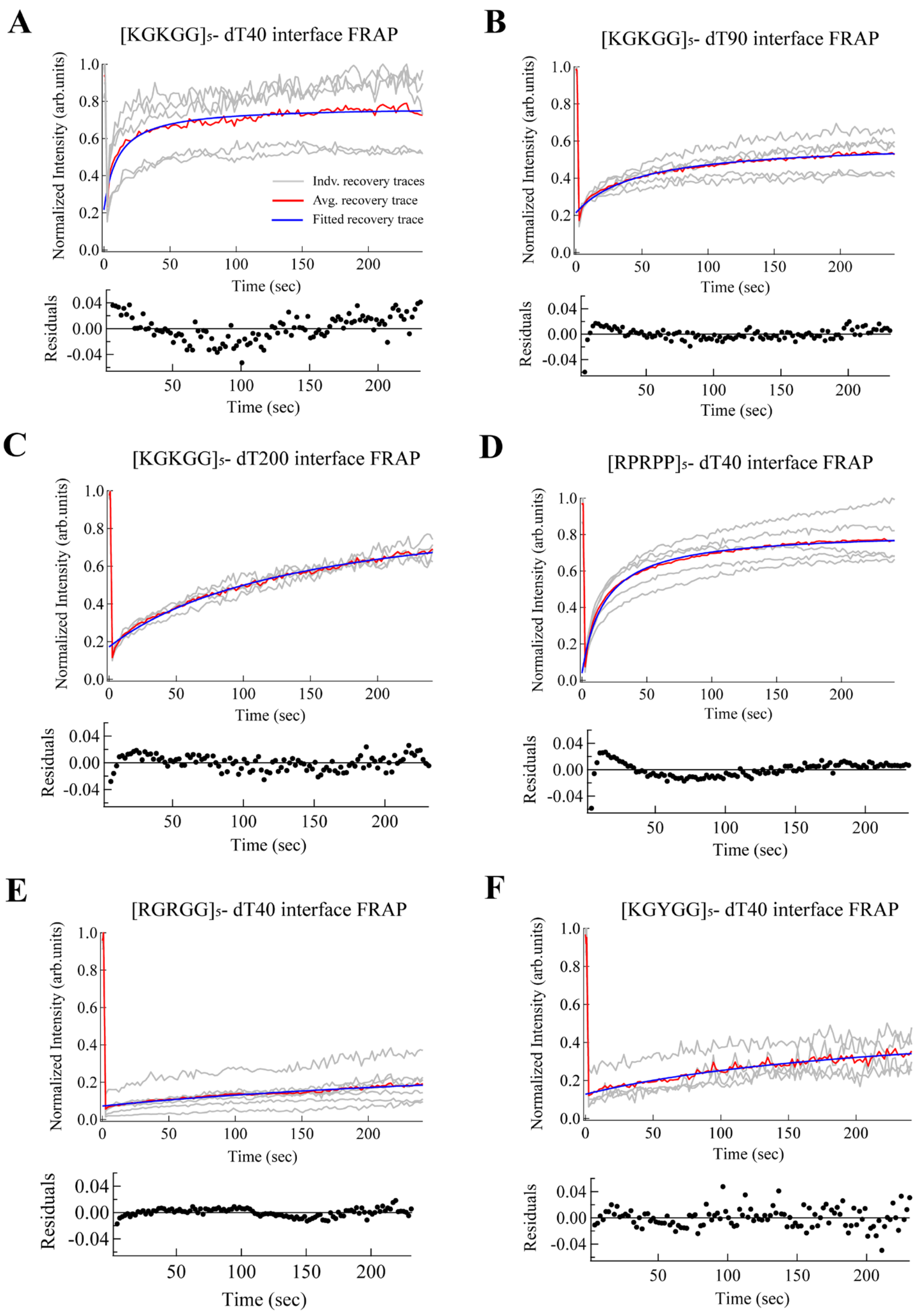


**Figure S11. Fitting of interfacial FRAP intensity traces for extracting recovery half-times and mobile fractions.** The top panel shows the normalized average recovery trace of $^{Hy647}$Tubulin (red), and the fitted recovery trace obtained using the function[8] $I(t) = a + b\ (t/\tau\ ½)/\ 1 + (t/\tau\ ½)$ (blue; see Methods, Table S1). The normalized average trace represents an average of 4-7 FRAP traces (grey) for each peptide-DNA combination across two independent sample replicates. The bottom panel shows the residuals (black solid circles). Data were analyzed for interfacial FRAP performed on $[KGKGG]_5$-dT40 condensates **(A)**; $[KGKGG]_5$-dT90 condensates **(B)**; $[KGKGG]_5$-dT200 condensates **(C)**; $[RPRPP]_5$-dT40 condensates **(D)**; $[RGRGG]_5$-dT40 condensates **(E)**; and $[KGYGG]_5$-dT40 condensates **(F).**

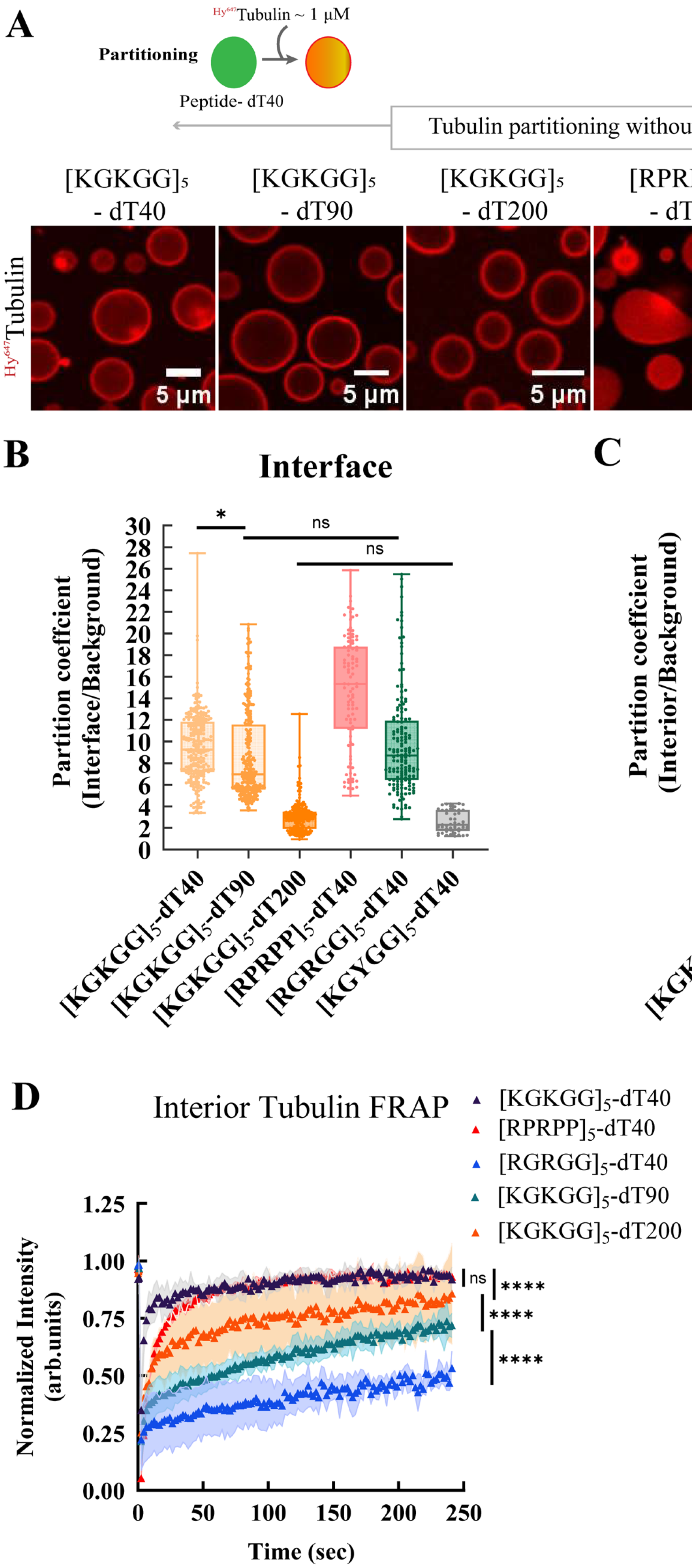
A
Hy647Tubulin ~ 1 μM
Partitioning
Peptide- dT40
Tubulin partitioning without GTP
[KGKGG]5 - dT40
[KGKGG]5 - dT90
[KGKGG]5 - dT200
[RPRPP]5 - dT40
[RGRGG]5 - dT40
[KGYGG]5 - dT40
Hy647Tubulin
5 μm
B
Interface
Partition coeffcient (Interface/Background)
ns
*
[KGKGG]5-dT40
[KGKGG]5-dT90
[KGKGG]5-dT200
[RPRPP]5-dT40
[RGRGG]5-dT40
[KGYGG]5-dT40
C
Interior
Partition coeffcient (Interior/Background)
D
Interior Tubulin FRAP
[KGKGG]5-dT40
[RPRPP]5-dT40
[RGRGG]5-dT40
[KGKGG]5-dT90
[KGKGG]5-dT200
Normalized Intensity (arb.units)
Time (sec)
ns
****

**Figure S12. Condensate material states regulate tubulin enrichment and mobility. (A)** A schematic and fluorescence microscopy images showing the recruitment behavior of tubulin into $[KGKGG]_5$-dT40, $[KGKGG]_5$-dT90, $[KGKGG]_5$-dT200, $[RPRPP]_5$-dT40, $[RGRGG]_5$-dT40, and $[KGYGG]_5$-dT40 condensates, as indicated, all containing 900 nM $^{Hy647}$Tubulin (red), 2.5 mg/mL peptide, 1.25 mg/mL dT40. Tubulin partition coefficient measurements at the interface **(B)** and interior **(C)** of condensates, as indicated. The individual data points represent measurements based on three independent replicates. Sample size: $[KGKGG]_5$-dT40 (Interface & Interior: n = 245), $[KGKGG]_5$-dT90 (Interface & Interior: n = 273), $[KGKGG]_5$-dT200 (Interface & Interior: n = 193), $[RPRPP]_5$-dT40 (Interface & Interior: n = 96), $[RGRGG]_5$-dT40 (Interface & Interior: n = 161), and $[KGYGG]_5$-dT40 (Interface & Interior: n = 48). The box represents the interquartile range from the 25$^{th}$ to the 75$^{th}$ percentile, with the middle line showing the median, the whiskers showing the minimum to maximum values, and the individual datapoints plotted as circular symbols. Statistical significance was determined using a one-way ANOVA Dunn's multiple comparisons test between the samples ("ns" means non-significant, * means $p < 0.05$, ** means $p < 0.01$, *** means $p < 0.001$, **** means $p < 0.0001$). For the interfacial partitioning, the associated *P* values are p = 0.021 for $[KGKGG]_5$-dT40 vs. $[KGKGG]_5$-dT90 condensates; p = 0.301 for $[KGKGG]_5$-dT40 vs. $[RGRGG]_5$-dT40 condensates; p = 0.366 for $[KGKGG]_5$-dT90 vs. $[RGRGG]_5$-dT40 condensates, $p > 0.99$ for $[KGKGG]_5$-dT200 vs. $[KGYGG]_5$-dT40 condensates, and $p < 0.0001$ for all other pairwise combinations. For the interior partitioning, the associated *P* values are p = 0.187 for $[KGKGG]_5$-dT90 vs. $[RGRGG]_5$-dT40 condensates, $p > 0.99$ for $[KGKGG]_5$-dT200 vs. $[KGYGG]_5$-dT40 condensates, and $p < 0.0001$ for all other pairwise combinations. **(D)** FRAP intensity-time traces of tubulin probe ($^{Hy647}$Tubulin) measured inside condensates as indicated. Sample conditions as described in (Fig 3E, F). Solid circles represent the average of 3-4 FRAP curves for each peptide-DNA combination across 2 independent replicates. Error bars in the curves represent one standard deviation (±1 s.d.). Statistical significance was determined using a one-way ANOVA Dunn's multiple comparisons test between the individual conditions, where "ns" means non-significant, **** means $P \leq 0.0001$.

**Table S1.** Extracted FRAP recovery half-times ($\tau_{1/2}$) and the goodness of fit ($\chi^2$) for the peptide-NA systems (corresponding to data shown in Figures 3F; 4E; and S11).

| | **[KGKGG]$_5$-dT40** | **[KGKGG]$_5$-dT90** | **[KGKGG]$_5$-dT200** | **[RPRPP]$_5$-dT40** | **[RGRGG]$_5$-dT40** | **[KGYGG]$_5$-dT40** |
|---|---|---|---|---|---|---|
| **$\tau_{1/2}$ (sec)** | 10.4 ± 1.2 | 47.2 ± 2.6 | 145.0 ± 7.8 | 17.5 ± 0.6 | 444.3 ± 88.4 | 247.9 ± 44.5 |
| **$\chi^2$** | 0.052 | 0.010 | 0.018 | 0.015 | 0.004 | 0.025 |

**Table S2.** Estimation of mobile fractions ($M_f$) from the FRAP data for the peptide-NA systems (corresponding to data shown in Figures 3F; 4E; and S11).

| | **[KGKGG]$_5$-dT40** | **[KGKGG]$_5$-dT90** | **[KGKGG]$_5$-dT200** | **[RPRPP]$_5$-dT40** | **[RGRGG]$_5$-dT40** | **[KGYGG]$_5$-dT40** |
|---|---|---|---|---|---|---|
| **$M_f$ (%)** | 71.2 ± 26.1 | 43.6 ± 13.3 | 65.5 ± 5.5 | 75.2 ± 15.5 | 13.7 ± 7.2 | 27.5 ± 12.8 |

### Supplementary Note 1:

#### Overview of stochastic model of microtubule nucleation and growth at condensate interfaces

We developed a coarse-grained stochastic model of microtubule (MT) nucleation, growth, and detachment at the surface of a spherical condensate droplet of radius $R$. The model treats tubulin as a continuous surface density field $n(\mathbf{u}, t)$ defined on a 2D grid mapped onto the droplet surface and represents each MT as a chain of discrete nodes in 3D space. The condensate interface viscosity $\eta_{\text{eff}}$ is the primary control parameter, encoding the density and cohesive strength of condensate–tubulin contacts at the droplet surface.

**Surface Tubulin adsorption**. Surface tubulin is assumed to adsorb from dilute phase according to Langmuir isotherm kinetics. The equilibrium surface density is

$$n_{\text{eq}} = n_{\text{max}} \frac{c_{\text{dilute}}}{K_d + c_{\text{dilute}}} \qquad (1)$$

where $n_{\text{max}}$ is the maximum surface saturation density, $c_{\text{dilute}}$ is the dilute phase tubulin concentration, and $K_d$ is the half-saturation constant.

**Surface dynamics.** The surface field evolves according to

$$\frac{\partial n}{\partial t} = k_{\text{ads}}(n_{\text{eq}} - n) + D_{\text{surf}} \nabla^2 n \qquad (2)$$

where the adsorption rate from dilute phase follows a Smoluchowski flux,

$$k_{\text{ads}} = \frac{k_0^{\text{ads}} D_{3\text{D}}}{R} \qquad (3)$$

and the 2D surface diffusivity obeys a Stokes–Einstein-like phenomenological relation,

$$D_{\text{surf}} = \frac{1}{\eta_{\text{eff}}} \qquad (4)$$

High interface viscosity, therefore, simultaneously slows lateral redistribution of surface tubulin and reduces the adsorption flux from dilute phase. The PDE is integrated numerically using an exact exponential relaxation for the adsorption term and an explicit finite-difference scheme for the diffusion term, with adaptive sub-steps to satisfy the stability condition $D_{\text{surf}} \Delta t / \Delta x^2 \leq 0.2$. MT dynamics are simulated as a continuous-time Gillespie (next-reaction) process with three classes of events: nucleation, growth, and tip detachment (peeling).

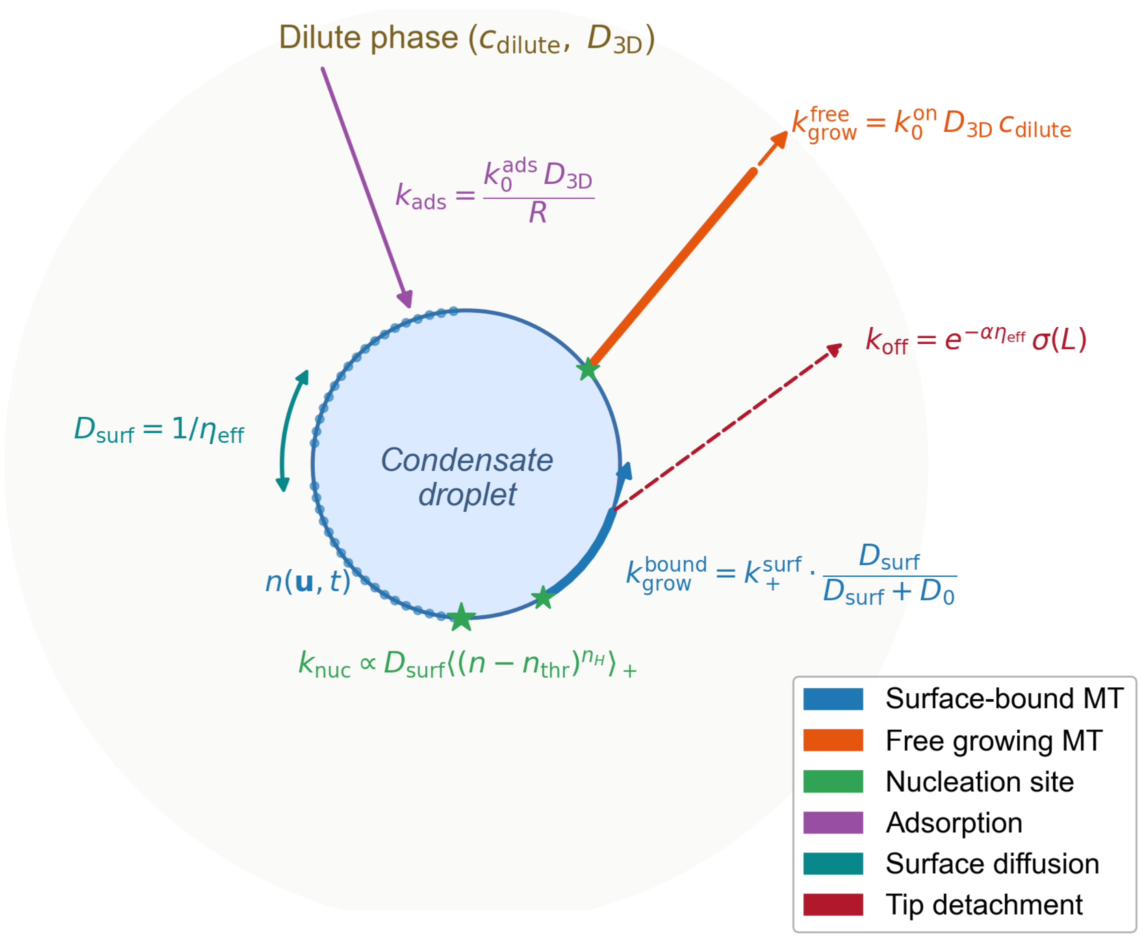


**Figure S13. Reaction scheme for the condensate surface-mediated MT growth model.** A spherical condensate droplet (blue, radius $R$) is surrounded by dilute solution containing free tubulin at concentration $c_{dilute}$ with 3D diffusivity $D_{3D}$. Tubulin adsorbs from dilute phase onto the droplet surface to form a 2D density field $n(\mathbf{u}, t)$ (blue dots) at rate $k_{ads} = k_0^{ads} D_{3D}/R$ (purple arrow), and redistributes laterally by surface diffusion with coefficient $D_{surf} = 1/\eta_{eff}$ (cyan double-headed arc), where $\eta_{eff}$ is the interface viscosity set by the condensate viscoelasticity. When local surface density exceeds a threshold, MT nucleation occurs (green star) at a rate proportional to $D_{surf}$. Surface-bound MTs (blue filament) grow along the droplet surface at rate $k_{grow}^{bound} = k_+^{surf} \cdot D_{surf}/(D_{surf} + D_0)$, which is directly throttled by interface viscosity through the diffusive-saturation factor (see Equation-6). Microtubule tip detachment occurs via escape at rate $k_{off}(L) = k_0^{off}\, e^{-\alpha\eta_{eff}}\, \sigma\left(\frac{L/R - \ell_{thresh}}{w_{thresh}}\right)$ (dashed red arrow; see Equation-10), where the exponential factor suppresses tip dissociation at high $\eta_{eff}$ and the sigmoidal gate $\sigma$ captures length-dependent bending stress at the contact zone. Once detached, free tips (orange filament) polymerize from dilute phase at rate $k_{grow}^{free} = k_0^{on} D_{3D} c_{dilute}$ (see Equation-7), independent of surface viscosity. The interplay between diffusion-limited surface growth and condensate material-dependent detachment gives rise to viscosity-dependent MT morphologies ranging from stalled surface-confined filaments (high $\eta_{eff}$, RGRGG-like) to radial asters (low $\eta_{eff}$, KGKGG-like). Filled circles mark MT anchor points fixed to the droplet surface.

**Nucleation.** The global nucleation rate is

$$k_{\text{nuc}} = \frac{k_0^{\text{nuc}}\, D_{\text{surf}}}{R^2} \sum_{i,j} \left\langle \left( \frac{n_{ij} - n_{\text{thresh}}}{n_{\text{thresh}}} \right)^{n_{\text{Hill}}} \right\rangle_+ \qquad (5)$$

where $\langle \cdot \rangle_+$ denotes the positive part, $n_{\text{thresh}}$ is the local density threshold required for productive nucleation, and $n_{\text{Hill}}$ is the cooperativity exponent. The $D_{\text{surf}}$ prefactor reflects that nucleation requires lateral assembly of a tubulin cluster, which is diffusion-limited. When a nucleation event fires, the anchor and initial tip are placed at the selected surface patch, and a fixed amount $\Delta n_{\text{nuc}}$ of tubulin is consumed locally.

**Growth for surface-bound tips.** The growth rate for a tip whose last node lies on the droplet surface is

$$k_{\text{grow}}^{\text{bound}} = k_+^{\text{surf}} \cdot \frac{D_{\text{surf}}}{D_{\text{surf}} + D_0} \qquad (6)$$

where $k_+^{\text{surf}}$ is the intrinsic on-rate and $D_0$ sets the crossover between diffusion-limited and reaction-limited regimes. This saturation form is the steady-state solution of a 2D diffusion-limited capture problem: the flux of tubulin delivered to a moving tip on the surface scales as $D_{\text{surf}}/(D_{\text{surf}} + D_0)$. High $\eta_{\text{eff}}$ reduces $D_{\text{surf}}$, directly throttling the growth rate. Each successful growth event additionally consumes $\Delta n_{\text{grow}}$ from the local surface patch; if the patch is depleted below this amount, the event is rejected (kinetic arrest).

**Growth for free tips.** Once a tip has detached from the surface, it polymerizes directly from dilute phase:

$$k_{\text{grow}}^{\text{free}} = k_0^{\text{on}}\, D_{\text{3D}}\, c_{\text{dilute}} \qquad (7)$$

Free tips are not throttled by surface viscosity.

**Simplified microtubule mechanics for free and bound tips**. At each growth step, the tip advances by a fixed arc length $\delta s$ along the surface (bound) or in 3D (free). The new direction is

$$\hat{d} = \hat{t} + \sigma\, \boldsymbol{\xi}, \qquad \boldsymbol{\xi} \sim \mathcal{N}(0, \mathbf{I}), \qquad (8)$$

normalized and for bound tips projected back onto the sphere. The angular noise amplitude for bound tips is set by worm-like chain statistics,

$$\sigma_{\text{bound}} = \sqrt{\delta s / L_p} \qquad (9)$$

where $L_p$ is the MT persistence length. Free tips use a fixed noise $\sigma_{\text{free}}$ that controls the angular spread of the aster.

**Tip detachment.** The rate at which a surface-bound tip detaches following Kramers-like escape dependence modified by a length-dependent force gate (Bell–Evans like model):

$$k_{\text{off}}(L) = k_0^{\text{off}}\, e^{-\alpha \eta_{\text{eff}}}\, \sigma\left( \frac{L/R - \ell_{\text{thresh}}}{w_{\text{thresh}}} \right) \qquad (10)$$

where $\sigma(\cdot)$ is the logistic function, $L$ is the current MT contour length, and $\ell_{\text{thresh}}$, $w_{\text{thresh}}$ set the length scale and width of the detachment gate. The factor $e^{-\eta_{\text{eff}}}$ captures the dissociation barrier, whereas α=1 is a phenomenological unit conversion factor that simply converts interfacial viscosity to adhesion free energy units. The denser, more cohesive interfaces (high $\eta_{\text{eff}}$, corresponding to RGRGG-type sequences) exponentially suppress tip peeling, while low-viscosity interfaces (KGKGG-type) allow spontaneous detachment. Long MTs accumulate bending stress at the contact zone, progressively lowering the effective barrier and triggering peeling.

### S2. Gillespie Simulation Algorithm

At each step of the Gillespie simulation loop, we do the following:

1. Compute all event rates: one nucleation rate $k_{\mathrm{nuc}}$; per-MT growth rates $k_{\mathrm{grow}}^{(i)}$ and detachment rates $k_{\mathrm{off}}^{(i)}$ for each MT $i$.
2. Draw the waiting time $\Delta t \sim \mathrm{Exp}(1/k_{\mathrm{tot}})$, where $k_{\mathrm{tot}} = \sum_i$ (all rates).
3. Advance the surface field by $\Delta t$ (adsorption + diffusion substeps).
4. Select and execute one event with probability proportional to its rate.
5. Record observables at pre-specified sample times.

Morphological observables reported are mean MT contour length $\langle L \rangle$, escape fraction $1 - f_{\mathrm{bound}}$ quantifying fraction of MTs whose tip has detached from the surface, and the aster order parameter

$$S = \langle \hat{t}_{\mathrm{tip}} \cdot \hat{n}_{\mathrm{tip}} \rangle, \qquad (11)$$

where $\hat{t}_{\mathrm{tip}}$ is the tip growth direction and $\hat{n}_{\mathrm{tip}}$ is the outward radial unit vector. $S \sim 1$ indicates a radial aster; $S \sim 0$ indicates tangential or tactoid-like organization.

### Table S3. Parameter Table

All simulations in this work used the parameter values listed below unless otherwise noted. Varied parameters ($\eta_{\mathrm{eff}}$, $c_{\mathrm{dilute}}$, $R$) are indicated in each figure panel.

| Symbol | Description | Value |
|---|---|---|
| **Primary** | | |
| $\eta_{\mathrm{eff}}$ | Interface viscosity (dimensionless) | varied (0.1–10) |
| $c_{dilute}$ | Dilute tubulin concentration (a.u.) | varied (2–20) |
| $R$ | Droplet radius (μm) | 8.0 |
| **Surface field** | | |
| $n_{\mathrm{max}}$ | Maximum surface density (a.u.) | 80 |
| $K_d$ | Langmuir half-saturation (a.u.) | 5.0 |
| $n_0/n_{\mathrm{eq}}$ | Initial surface loading fraction | 0.45 |
| $N_x \times N_y$ | Surface grid resolution | 40 × 40 |
| $L_x \times L_y$ | Grid physical size (μm) | 20 × 20 |
| **Transport** | | |
| $D_{\mathrm{3D}}$ | Dilute phase diffusivity (a.u.) | 1.0 |
| $k_0^{\mathrm{ads}}$ | Adsorption prefactor | 0.14 / 8.0 |
| $k_0^{\mathrm{on}}$ | Free-tip on-rate prefactor | $3 \times 10^{-3}$ |
| **Nucleation** | | |

| Symbol | Description | Value |
|---|---|---|
| $k_0^{\mathrm{nuc}}$ | Nucleation prefactor | $2 \times 10^{-4}$ |
| $n_{\mathrm{thresh}}$ | Surface density nucleation threshold | 6.0 |
| $n_{\mathrm{Hill}}$ | Hill cooperativity exponent | 3.0 |
| $\Delta n_{\mathrm{nuc}}$ | Tubulin consumed per nucleation event | 8.0 |
| **Growth** | | |
| $k_+^{\mathrm{surf}}$ | Surface on rate ($s^{-1}$) | 0.11 |
| $D_0$ | Diffusion-saturation crossover scale | 4.0 |
| $\Delta n_{\mathrm{grow}}$ | Tubulin consumed per growth step | 5.0 |
| $\delta s$ | Growth step length (μm) | 0.18 |
| $L_p$ | MT persistence length (μm) | 1000 |
| $\sigma_{\mathrm{free}}$ | Angular noise for free tips (rad) | 0.05 |
| **Tip detachment** | | |
| $k_0^{\mathrm{off}}$ | Basal detachment rate ($s^{-1}$) | 0.005 |
| $\ell_{\mathrm{thresh}}$ | Length gate threshold ($L/R$) | 0.30 |
| $w_{\mathrm{thresh}}$ | Length gate width ($L/R$) | 0.30 |
| **Run limits** | | |
| $N_{\mathrm{MT}}^{\max}$ | Maximum MTs per droplet | 28 |
| $T_{\mathrm{end}}$ | Simulation duration (s) | 2000 |

### Supplementary Video Legend

#### Video S1

Addition of 10 µM tubulin doped with 900 nM $^{Hy647}$Tubulin plus 2 mM GTP, to $[KGKGG]_5$–dT40 condensates doped with 500 nM $^{AF488}[KGKGG]_5$, visualized as an overlay of $^{Hy647}$Tubulin & $^{AF488}[KGKGG]_5$ fluorescence, showing condensate dissolution and MT nucleation and growth. The composition of the condensate system is 2.5 mg/mL $[KGKGG]_5$ and 1.25 mg/mL dT40, with final sample containing 60 mM PIPES (pH 6.9), 1.5 mM $MgCl_2$, 0.38 mM EGTA and 12 mM DTT.

#### Video S2

Addition of 5 µM tubulin doped with 500 nM $^{Hy647}$Tubulin plus 1 mM GTP, to $[KGKGG]_5$–dT40 condensates doped with 500 nM $^{AF488}[KGKGG]_5$, visualized as an overlay of $^{Hy647}$Tubulin & $^{AF488}[KGKGG]_5$ fluorescence, showing MT nucleation and growth at the interface of condensate. The condensate composition and sample conditions are as reported in Video S1. Scale bar is 5 µm.

#### Video S3

Addition of tubulin buffer plus 1 mM GTP to $[KGKGG]_5$–dT40 condensates doped with 500 nM $^{AF488}[KGKGG]_5$, visualized from $^{AF488}[KGKGG]_5$ fluorescence, showing intact condensates and absence of MT growth. The condensate composition and sample conditions are as reported in Videos S1 and S2. Scale bar is 5 µm.

#### Video S4

Addition of 10 µM tubulin doped with 900 nM $^{Hy647}$Tubulin plus 2 mM GTP, to $[KGKGG]_5$–dT40 condensates doped with 500 nM $^{AF488}[KGKGG]_5$, visualized as an overlay of $^{Hy647}$Tubulin & $^{AF488}[KGKGG]_5$ fluorescence, showing rapid MT nucleation and growth from potential $[KGKGG]_5$–dT40 nano-scale clusters. The composition of the condensate system used here is 2.5 mg/mL $[KGKGG]_5$ and 5.25 mg/mL dT40, with the final sample containing 60 mM PIPES (pH 6.9), 1.5 mM $MgCl_2$, 0.38 mM EGTA, and 12 mM DTT. Scale bar is 5 µm.

#### Video S5

Addition of 10 µM tubulin doped with 900 nM $^{Hy647}$Tubulin plus 2 mM GTP, to $[RGRGG]_5$–dT40 condensates doped with 500 nM $^{AF488}[RGRGG]_5$, visualized as an overlay of $^{Hy647}$Tubulin & $^{AF488}[RGRGG]_5$ fluorescence, showing stable condensates and short MT asters. The condensate composition and sample conditions are as reported in Videos S1 and S2. Scale bar is 5 µm.

#### Video S6

Representative FRAP of $^{Hy647}$Tubulin at the interface of $[KGKGG]_5$–dT40 condensates. The composition of the condensate system is 2.5 mg/mL $[KGKGG]_5$ and 1.25 mg/mL dT40, with the final sample containing 60 mM PIPES (pH 6.9), 1.5 mM $MgCl_2$, 0.38 mM EGTA, and 12 mM DTT. The concentration of the labelled component is 900 nM. Measurements were performed 15 min after sample preparation. Scale bar, 2 μm.

**Video S7**

Representative FRAP of $^{Hy647}$Tubulin in the interior of $[KGKGG]_5$–dT40 condensates. The sample composition and conditions are as described in Video S6; Figs 3F and 4E. Sample analysis and imaging were done 15 min after preparation. Scale bar, 2 μm.

**Video S8**

Representative FRAP of $^{Hy647}$Tubulin at the interface of $[RGRGG]_5$–dT40 condensates. The sample composition and conditions are as described in Video S6; Figs 3F and 4E. Scale bar, 2 μm.

**Video S9**

Representative FRAP of $^{Hy647}$Tubulin in the interior of $[RGRGG]_5$–dT40 condensates. The sample composition and conditions are as described in Video S6; Figs 3F; and 4E. Scale bar, 1 μm.

**Video S10**

Representative FRAP of $^{Hy647}$Tubulin at the interface of $[RPRPP]_5$–dT40 condensates. The sample composition and conditions are as described in Video S6; Figs 3F and 4E. Scale bar, 2 μm.

**Video S11**

Representative FRAP of $^{Hy647}$Tubulin in the interior of $[RPRPP]_5$–dT40 condensates. The sample composition and conditions are as described in Video S6; Figs 3F and 4E. Scale bar, 2 μm.

**Video S12**

Representative FRAP of $^{Hy647}$Tubulin at the interface of $[KGKGG]_5$–dT90 condensates. The sample composition and conditions are as described in Video S6; Figs 3F and 4E. Scale bar, 2 μm.

**Video S13**

Representative FRAP of $^{Hy647}$Tubulin in the interior of $[KGKGG]_5$–dT90 condensates. The sample composition and conditions are as described in Video S6; Figs 3F and 4E. Scale bar, 2 μm.

**Video S14**

Representative FRAP of $^{Hy647}$Tubulin at the interface of $[KGKGG]_5$–dT200 condensates. The sample composition and conditions are as described in Video S6; Figs 3F and 4E. Scale bar, 2 μm.

**Video S15**

Representative FRAP of $^{Hy647}$Tubulin in the interior of $[KGKGG]_5$–dT200 condensates. The sample composition and conditions are as described in Video S6; Figs 3F and 4E. Scale bar, 2 μm.

**Video S16**

Representative FRAP of $^{Hy647}$Tubulin at the interface of $[KGYGG]_5$–dT40 condensates. The sample composition and conditions are as described in Video S6; Figs 3F and 4E. Scale bar, 2 μm.

**Video S17**

MT assembly at a high-viscosity condensate interface ($\eta_{eff} = 10$). Time-lapse of a stochastic simulation showing tubulin adsorption, lateral surface diffusion, nucleation, and MT growth on a spherical condensate. Slow surface diffusion ($D_{surf} \lesssim D_0$) puts the system in the diffusion-limited regime yielding sparser, shorter MTs despite identical bulk tubulin. Direct comparison with Video S18 illustrates that interface viscosity, not bulk tubulin availability, controls assembly under these conditions.

**Video S18**

MT assembly at a low-viscosity condensate interface ($\eta_{eff} = 0.1$). Time-lapse of a stochastic simulation showing tubulin adsorption, lateral surface diffusion, nucleation, and MT growth on a spherical condensate. Fast surface diffusion ($D_{surf} \gg D_0$) puts the system in the reaction-limited regime: nucleation is efficient and bound tips grow close to the intrinsic on-rate.
